# Two highly selected mutations in the tandemly duplicated *CYP6P4a* and *CYP6P4b* drive pyrethroid resistance in *Anopheles funestus*

**DOI:** 10.1101/2024.03.26.586794

**Authors:** Nelly M.T. Tatchou-Nebangwa, Leon M. J. Mugenzi, Abdullahi Muhammad, Derrick N. Nebangwa, Mersimine F.M. Kouamo, Carlos S. D. Tagne, Theofelix A. Tekoh, Magellan Tchouakui, Stephen M. Ghogomu, Sulaiman S. Ibrahim, Charles S. Wondji

## Abstract

Gaining a comprehensive understanding of the genetic mechanisms underlying insecticide resistance in malaria vectors is crucial for optimising the effectiveness of insecticide-based vector control methods and developing diagnostic tools for resistance management. Considering the heterogeneity of metabolic resistance in major malaria vectors, the implementation of tailored resistance management strategies is essential for successful vector control. In this study, we provide evidence demonstrating that two highly selected mutations in the tandemly duplicated cytochrome P450 genes namely *CYP6P4a* and *CYP6P4b*, are driving pyrethroid insecticide resistance in the major malaria vector *Anopheles funestus,* in West Africa. Through a continent-wide polymorphism survey, we observed heightened indications of directional selection in both genes between 2014 and 2021. By conducting *in vitro* insecticide metabolism assays with recombinant enzymes expressed from both genes, we established that mutant alleles under selection exhibit higher metabolic efficiency compared to their wild-type counterparts. Furthermore, using the GAL4-UAS transgenic system, we demonstrated that transgenic *Drosophila melanogaster* flies overexpressing mutant alleles displayed an increased resistance to pyrethroids. These findings were in agreement with *in silico* characterisation, which highlighted changes in enzyme active site architecture that enhance the affinity of mutant alleles for type I and II pyrethroids. Furthermore, we developed two DNA-based assays capable of detecting the CYP6P4a-M220I and CYP6P4b-D284E mutations, showing their current confinement to West Africa. Genotype/phenotype correlation analyses revealed that these markers are strongly associated with resistance to types I and II pyrethroids and combine to drastically reduce the efficacy of pyrethroid bednets. Overall, our study makes available two field-applicable insecticide resistance molecular markers that will help in the monitoring and better management of insecticide resistance in West Africa.

**Teaser:** Two field-applicable diagnostic tools for detecting metabolic resistance in *Anopheles funestus* to enhance insecticide resistance management in West Africa.

## Introduction

Globally, malaria continues to be a leading cause of human mortality, disproportionately affecting sub-Saharan Africa. In 2022, an estimated 249 million cases were reported, resulting in 608 thousand deaths, with more than 90% occurring in sub-Saharan Africa and in children under the age of 5 ^1^. The control of the disease heavily relies on the use of insecticide-based vector control tools, primarily long-lasting insecticide-treated bed nets (LLINs) and indoor residual sprays (IRS)^2^ which have achieved an unprecedented reduction in the disease burden, accounting for about 78% of the decline in malaria cases between 2000 and 2015 ^3,4^. Unfortunately, the massive use of insecticides has exerted a selective pressure on mosquito populations, leading to the emergence of resistance mechanisms that enable mosquitoes to survive exposure to these chemicals. Resistance to all major insecticide classes currently used in public health including pyrethroids, organochlorines, carbamates, and organophosphates, has been extensively reported in the dominant African malaria vectors ^5^. Among these, resistance to pyrethroids is of particular concern as pyrethroids are the primary class of insecticides recommended by the WHO for bed net impregnation ^1^. Notably, the upsurgence of malaria in the Kwazulu/Nata province in South Africa between 1996 and 2001 was attributed to the emergence of pyrethroid resistance in the major malaria vector *Anopheles funestus* Giles ^6^. Therefore, the dissemination of the genetic basis of insecticide resistance and the development of tools for early detection and anticipation of resistance remain pivotal in devising effective resistance management strategies to control malaria vectors ^7,8^. The main mechanisms of insecticide resistance in malaria vectors are target-site resistance and metabolic resistance ^9,10^. While significant progress has been made in understanding target-site resistance, with key mutations identified over 25 years ago ^11^, unravelling the complexities of metabolic resistance has proven more challenging. This complexity arises from the involvement of multiple gene families, encompassing dozens of genes, with an estimated total of around 200 genes potentially contributing to the resistance phenotype ^9,10^. Furthermore, even after identifying candidate resistance genes, numerous resistance pathways exist, including mutations in regulatory and coding regions, transposable elements ^12^. Consequently, detecting genetic variants responsible for metabolic resistance and establishing diagnostic tools though pivotal, are challenging. Nonetheless, significant progress has been made with the detection of the first DNA-based maker in the GST gene *GSTe2* conferring DDT resistance ^13^ in *An. funestus*. Similarly, the elucidation of cis-regulatory elements associated with pyrethroid resistance in southern Africa led to the development of the first DNA-based markers for cytochrome P450-mediated metabolic resistance in *An. funestus* in 2019 ^14,15^. However, it is important to note that metabolic resistance in Africa exhibits heterogeneity, with different genes driving resistance in various regions due to barriers to gene flow, emphasising the necessity of tailored resistance management strategies ^16–18^. In this context, two tandemly duplicated cytochrome P450 genes, *CYP6P4a* and *CYP6P4b,* have been consistently shown to be the most over-expressed genes in pyrethroid resistant populations of *An. funestus* in West Africa ^15^. However, their direct involvement in resistance and associated variants have not been identified, hindering the detection and monitoring of resistance in these populations. In this study, we conducted an analysis of the Africa-wide polymorphism of the duplicated *CYP6P4a* and *CYP6P4b* genes, revealing heightened signatures of directional selection of predominant alleles between 2014 and 2021. Through *in silico* modelling and molecular docking, we predicted a greater affinity of the selected alleles to pyrethroids. Moreover, *in vitro* insecticide metabolism assays and *in vivo* transgenic expression in *Drosophila melanogaster* flies followed by insecticide-contact bioassays demonstrated that the selected alleles of both genes exhibited higher ability to metabolise both type I and II pyrethroids, resulting in significantly increased survivorship compared to the wild-type alleles. In addition, we have developed two DNA-based resistance diagnostic tools that will allow the spatio-temporal tracking of the spread of resistance across the continent. Currently, these markers have been detected in West African countries emphasising the need for routine monitoring to limit the spread of the resistance genes in other regions. We also show here that both CYP6P4a-M220I and CYP6P4b-D284E mutations are robust markers that predict resistance status co-dominantly, revealing a significant reduced efficacy of pyrethroid-only bed nets.

## Results

### Africa-wide polymorphism analysis of *CYP6P4a* and *CYP6P4b* reveals signatures of directional selection

To assess the genetic diversity of resistant *An. funestus* mosquitoes in different regions of Africa, and to detect variants associated with pyrethroid resistance, we carried out polymorphism analysis of the full-length coding region of *CYP6P4a* (1533 bp) and *CYP6P4b* (1542 bp). Mosquito samples involved in the study were collected in 2021 from representative countries of the four Sub-Saharan regions of African: Southern (Malawi and Mozambique), Eastern (Uganda), Central (Cameroon) and Western (Ghana and Benin). Polymorphism patterns showed a high level of homogeneity and reduced diversity within individual populations but with significant variation between geographical regions (Table S1). The *CYP6P4a* gene had a total of 17 haplotypes with 66 polymorphic sites, including 9 non-synonymous (Table S1). The resistant Ghanaian mosquito population showed reduced haplotype diversity (hd = 0.583), but with a high number of polymorphic sites; this rather due to a singleton that differed significantly from the predominant haplotype (6/9). Amino acid phylogenetic tree analysis revealed that the majority of Ghana sequences clustered together and were distinct from other populations (Fig. 1A). The haplotype network also indicated the presence of a major haplotype (H1) (Fig. 1B) circulating in the population, with 6 out of 9 sequences bearing it (Fig. 1C). Benin sequences also showed the existence of a major haplotype (5/6) that clustered away from the Ghanian population although both are in West Africa (Fig. 1B). Similarly, *CYP6P4b* demonstrated a reduction in diversity across all resistant populations. A total of 24 haplotypes were identified, encompassing 69 polymorphic sites of which 16 were non-synonymous mutations (Table S1). Again, the Ghanaian population displayed a significant reduction in diversity, characterised by two non-synonymous mutations, a low haplotype diversity (Hd) of 0.216, and a low nucleotide diversity (π) of 0.00014 (Table S1). Phylogenetic analyses confirmed the distinct clustering of all Ghanaian sequences from other populations, (Fig. 1D) and the prevalence of a predominant haplotype (H1) present among 16 out of 18 sequences (Figs. 1E). For both genes, the southern populations displayed no diversity (Hd = 0, D = 0, π = 0) (Table S1), with a single allele (Figs. 1A and 1D) and a single haplotype (H3) (Figs. 1B and 1E). Also, the Cameroon and Uganda sampled populations displayed no diversity in *CYP6P4a* (Hd = 0, D = 0, π = 0) and indications of selection of the *CYP6P4b* gene (Table S1), as most sequences from these regions grouped together with a common major haplotype (H2) (Figs. 1B and 1D). These haplotypes were fixed in the sampled population in Uganda (CYP6P4a = 10/10, CYP6P4b = 10/10) and close to fixation in the sampled population of Cameroon (CYP6P4a = 12/12, CYP6P4b = 5/7) (Figs. 1C and S2B B). Contrastingly, the insecticide-susceptible FANG population exhibited the highest diversity (hd = 1 for *CYP6P4a* and hd = 0.855 for *CYP6P4b*) with no signs of selection. This suggests an association between the selective sweep observed in the four regions and pyrethroid resistance.

**Figure 1:**
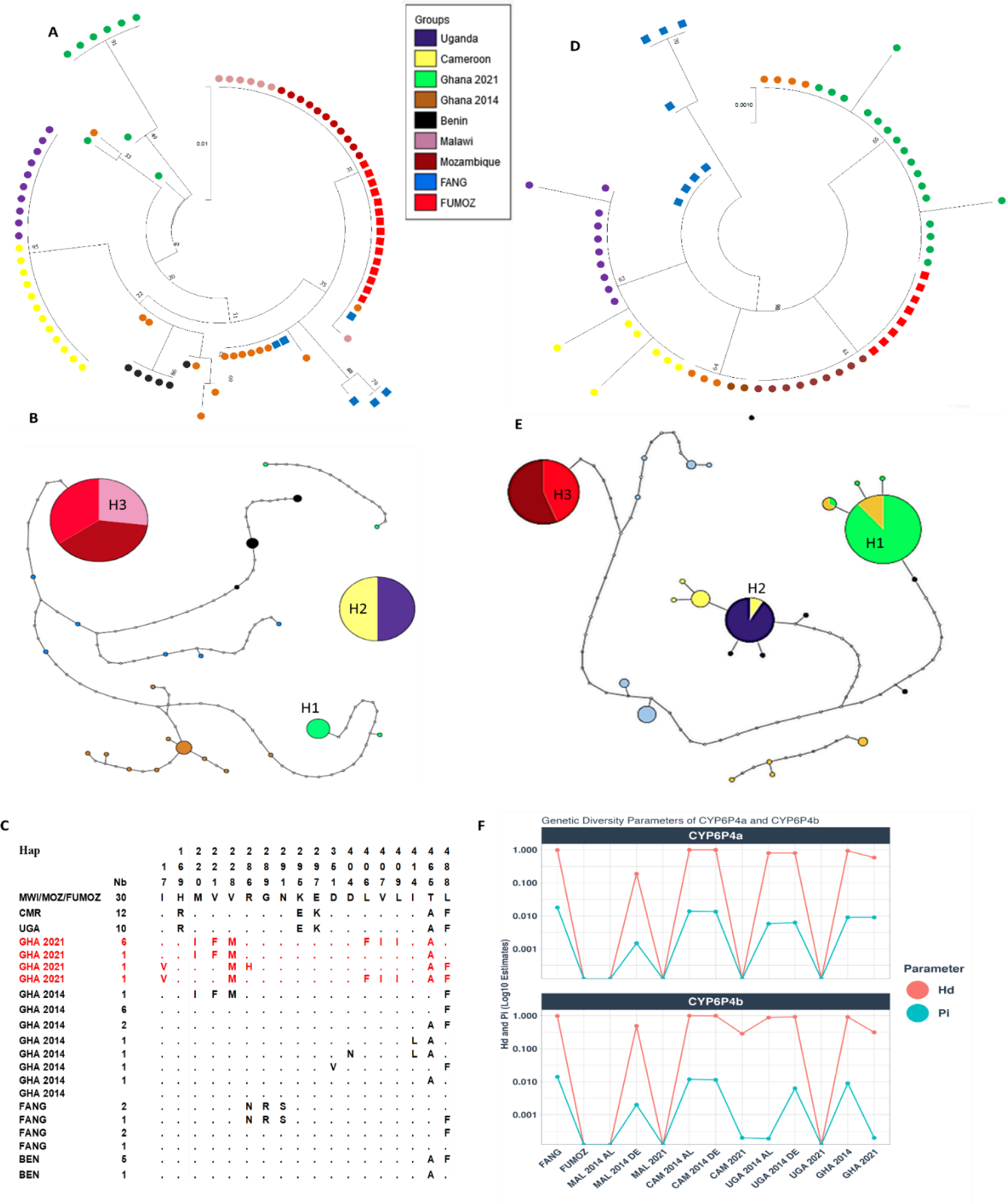
Schematic representation of haplotypes and genetic diversity of CYP6P4a and CYP6P4b in resistant mosquitoes, FUMOZ and FANG. (A) Phylogenetic tree for CYP6P4a plotted using the JTT model, showing selection of major alleles in all regions in 2021; (B) Haplotype network for CYP6P4a revealing a fixed haplotype common Southern Africa, a fixed haplotype common to Uganda and Cameroon, and major haplotypes in Ghana and Benin, but high diversity in FANG; (C), Schematic representation of haplotypes of CYP6P4a with mutations occurring in Ghana highlighted in red; (D) Phylogenetic tree for CYP6P4b plotted using the JTT model, showing selection of major alleles in all regions in 2021; (E), CYP6P4b haplotype network. A fixed haplotype is observed in Southern population (Malawi and FUMOZ) and the formation of a major haplotype common to Uganda and Cameroon; (F) Plot of genetic diversity parameters of CYP6P4a and CYP6P4b across Africa showing the signatures of a directional selection of CYP6P4a and CYP6P4b between 2014 and 2021 in haplotype diversity (Hd) and nucleotide diversity (π).

### Africa-wide temporal evolution of *CYP6P4a* and *CYP6P4b* is associated with increase in pyrethroid resistance

Capturing the evolutionary pattern of the genetic variation of genes is important in the assessment of the impact of insecticide pressure on the population structure. For this aim, we performed a temporal comparison of the diversity of the genes between 2014 and 2021. A striking reduction in genetic diversity is observable in both genes in the populations of Uganda, Cameroon, and Ghana. (Fig. 1F). For *CYP6P4a,* an overall drastic reduced diversity between 2014 and 2021 in Ghana, Cameroon, and Uganda is noted, while Malawi already exhibited signatures of selection by 2014. In Ghana, the temporal selection was evident through a reduction in haplotype diversity from 0.934 (h = 11) in 2014 to 0.583 (h = 4) in 2021 with the selection of a predominant allele in the population. An even stronger signature of selection was observed in Cameroon [(2014: D = 2.81, P < 0.01; Hd = 0.99; π = 0.01) vs (2021: D = 0, Hd = 0, π = 0)] and Uganda [(2014: D = 2.25, P < 0.05; Hd = 0.78; π = 0.01) vs. (2021: D = 0; Hd = 0, π = 0)] (Table S1and S2, and Fig. S1A and S1B). Both populations now sharing a common major haplotype. In contrast, the Malawian population exhibited low diversity in both 2014 (D = −1.80345; Hd = 0.097; π = 0.00076) and 2021 (D = 0; Hd = 0.0; π = 0), which can be attributed to earlier directional selection observed in the nearby genes *CYP6P9a/b* since 2010 ^19^ (Tables S1 and S2 and Fig. S1A and S1B). The same pattern was observed for *CYP6P4b*, where the genetic diversity was high in 2014 with no indications of directional selection in either Ghana (D = 2.24, P <0.05; Hd = 0.917; π = 0.009), Uganda (D = 2.09, P< 0.05; Hd = 0.89; π = 0.00592), or Cameroon (D = 2.2, P< 0.05, Hd = 0.994, π = 0.012) (Table S2 and Figs, S1 D, S1 E, and S1 F). The Malawi population however presented significant reduced diversity, with the gene undergoing selection (D = −1.84934; Hd = 0.273, π = 0.005) (Table S2 and Fig. 1F). In 2021, a marked reduction in gene diversity was observed in the same populations, with signs of strong selection: Ghana (D = −1.508; Hd = 0.216; π = 0.00014), Cameroon (D = −1.0062; Hd = 0.286; π = 0.00019) (Table S1 and Fig. S1 C). The Uganda and Malawi populations exhibited no diversity in 2021, with a unique allele in the population that had undergone selection to fixation prior to 2014 (Table S1, Figs. 1C, and S1 B).

### Africa-wide coding sequence polymorphisms of *CYP6P4a* and *CYP6P4b*

Distinctive mutations specific to each resistant population were identified through comparison of sequences from selected alleles in resistant mosquito populations collected in 2021 and sequences from the susceptible lab strain FANG. For *CYP6P4a*, the dominant allele in Ghana contained six mutations: *6P4a-I*^220^*F*^221^*M*^228^*F*^406^*I*^407^*I*^409^ hereby also referred to as 6P4a-GHA. The major allele common to Cameroon and Uganda had three mutations: *6P4a-R*^169^*E*^295^*K*^297^ and for FANG, one of the dominant alleles having three mutations: *6P4a-N*^286^*R*^289^*S*^291^, hereby also referred to as 6P4a-FANG. All the southern populations had a single *6P4a-L*^488^ allele circulating in the population (Fig. 1C). Concerning *CYP6P4b*, the dominant allele in Ghana had a single characteristic mutation: *6P4b-E*^284^, hereby also referred to as 6P4b-GHA, while the dominant allele common to Uganda and Cameroon had the *6P4b-N*^288^ mutation. The FANG allele used for downstream analyses was also identified as *6P4b-T*^291^*V*^294^*Y*^399^, hereby also referred to as 6P4b-FANG (Fig. S2B B). Again, the southern Africa populations had the same *6P4b-H*^26^ allele. These mutations were mapped to important domains of cytochrome P450s (Figs. S2A and S2B A) and the dominant alleles were used for downstream functional genomic validations.

### Mutations in *CYP6P4a* increase enzyme’s affinity for pyrethroids

To predict the impact of detected polymorphisms on the enzyme structure and enzyme-insecticide interactions, we performed computational modelling of CYP6P4a and molecular docking of pyrethroid ligands within the active sites of modelled enzymes (Fig. S3 K). All protein structures were modelled using Alphafold2, which significantly enhances the precision of structure prediction based on innovative neural network architecture ^20^. Template multiple sequence alignments (MSA) for Ghana *6P4a-I*^220^*F*^221^*M*^228^*F*^406^*I*^407^*I*^409^ (mutant) and FANG *6P4a-N*^286^*R*^289^*S*^291^ (wild-type) exhibited good sequence depth and coverage, with >30 template sequences as required for accurate Alphafold modelling (Fig. S3 E and F). All 3-D atomic resolution structural coordinates were calculated with high accuracy, reflected in the high average predicted Local Distance Difference Test score (pLDDT > 90%) of the modelled structures (Figs. S3 A and S3 C). The pLDDT score estimates the prediction confidence per residue site. Model confidence was further confirmed by the low average predicted aligned error (PAE < 5Å) that measures the distance error between corresponding estimated and aligned template residues (Figs. S3 B and S3 D). Structural superposition of wild-type and mutant alleles produced a mean backbone accuracy of 0.37Å r.m.s.d (Cα root mean square deviation at 95% residue coverage, confidence interval = 0.01– 3.16Å) (Figs. S3G and S3H). The highest backbone deviation occurred in the *K*^280^ *– M*^300^ helix-loop domain with a mean r.m.s.d. of 1.41Å, perhaps, strongly indicative of resulting conformational change elicited by three (*M*^220^*I, V*^221^*F, V*^228^*M*) of the six 6P4a-GHA mutations localised on the third N-terminal helix adjacent to this *K*^280^ *– M*^300^ domain in both models. As a comparison point, the approximate diameter of a carbon atom is 1.5Å, which is roughly equal to the 1.41Å r.m.s.d. displacement of the helix-loop. The heme molecule binding mode from the *CYP3A4* template crystal structure was mapped onto 6P4a-GHA and 6P4a-FANG models prior to docking, producing a mean accuracy of 0.35Å r.m.s.d. (All atom root-mean-square deviation between crystal and computationally predicted heme binding modes) (Fig. S3 J).

Following structural modelling, molecular docking was carried out to assess binding parameters of the pyrethroid ligands and protein models. Overall, the 3-D modelling of CYP6P4a–substrate interactions exhibited good binding scores and limited clashes in the active site. Ligand docking to 6P4a-GHA and 6P4a-FANG revealed contrasting binding conformations and parameter. Docked poses with functional sites approaching above the heme catalytic iron centre within 1.5 - 6.5Å to account for optimal van der Waals clashes were identified. The most enriched pyrethroid binding modes involved the 4’-phenoxybenzyl moiety, than any other moiety, approaching above the heme iron within 1.5 - 6.5Å and were thus, categorised herein as productive poses (Fig. 2A-2D). Subsequent analyses were performed on productive poses for each pyrethroid bound to 6P4a-FANG or 6P4a-GHA. The differences in the average distances between the 4’-phenoxy spot of all deltamethrin binding modes and the heme iron of the 6P4a-GHA (3.2 ± 0.42Å) and 6P4a-FANG (4.7 ± 0.38Å) models, as well as permethrin productive poses bound to 6P4a-GHA (3.4 ± 0.6Å) and 6P4a-FANG (4.3 ± 0.5Å) models were significant (p < 0.018) but modest, however, both pyrethroids exhibited shorter and more favorable distances for ring hydroxylation at the 4’ position when bound to the 6P4a-GHA model compared to 6P4a-FANG (Fig. 2A-2D, Table S3). Thermodynamic evaluation of affinity of binding for the selected poses predicted that 6P4a-GHA had consistent and significantly (p < 0.038) greater affinity for deltamethrin (MM affinity −9.04 ± 0.11 Kcal/mol; GBVI/WSA ΔG −35.5 ± 0.68 Kcal/mol; and S score −7.4 ± 0.03) than 6P4a-FANG (MM affinity −8.6 ± 0.13 Kcal/mol; GBVI/WSA ΔG −34.2 ± 1.3 Kcal/mol; and S. score −7.1 ± 0.04) (Table S3). Likewise, significantly (p < 0.029) better binding affinity for permethrin was predicted for the 6P4a-GHA enzyme (MM affinity −8.85 ± 0.28 Kcal/mol; GBVI/WSA ΔG −34.8 ± 1.2 Kcal/mol; S score −7.1 ± 0.05) compared to 6P4a-FANG (MM affinity −8.17 ± 0.11 Kcal/mol; GBVI/WSA ΔG −33.5 ± 0.9 Kcal/mol; and S score −6.8 ± 0.05) (Table S3).

**Figure 2:**
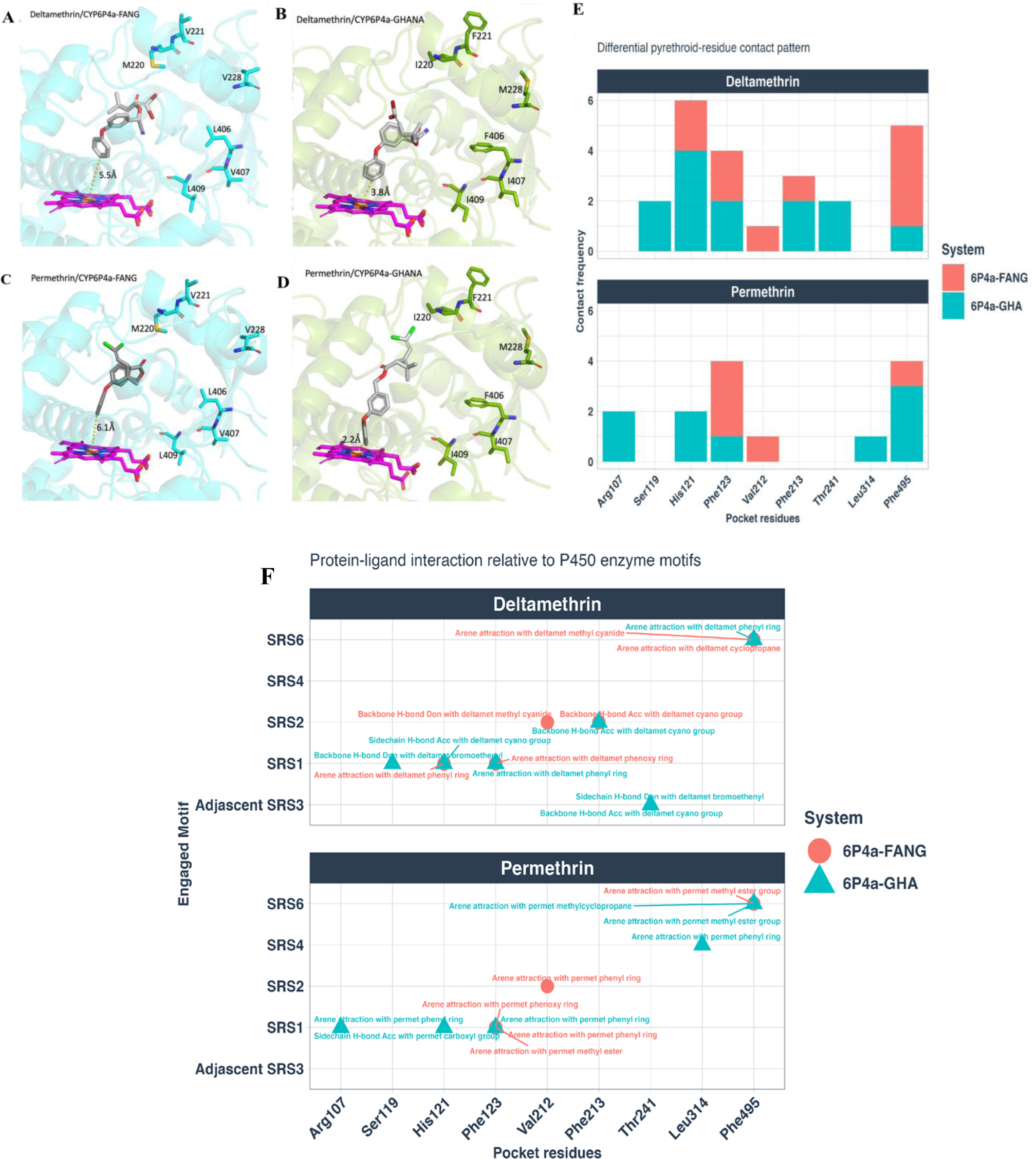
Modeling the CYP6P4a active site and impact of Amino Acid changes on pyrethroid Binding. Predicted 3-D binding modes of representative pyrethroid poses with 4’-phenoxy spot approaching above the heme iron at a distance ranging between 1.5 - 6.5Å. Deltamethrin bound to **(A)** CYP6P4a-FANG, **(B)** CYP6P4a-Ghana; and Permethrin bound to (**C).** CYP6P4a-FANG, **(D),** CYP6P4a-Ghana. models. Permethrin and Deltamethrin represented in stick format (colored grey), CYP6P4a-FANG (colored cyan) and CYP6P4a-Ghana (colored green) protein models are represented in cartoon with protruded wildtype (CYP6P4a-FANG) vs. mutant (CYP6P4a-Ghana) residues shown as sticks. Heme atoms also in sticks and colored purple. Distances between possible sites of metabolism and heme iron are annotated in Angstrom. **Protein-ligand interaction fingerprint between pyrethroid and protein models**. **(E),** Differential pyrethroid-residue contact pattern for deltamethrin (top panel) and permethrin (bottom panel) interacting with both wilt-type and mutant protein models. **(F)**, Interactions portray substrate recognition site (SRS) residues and motifs of wild- and mutant-type proteins interacting with pyrethroids; deltamethrin (first panel) and permethrin (second panel). Comparing the 6P4a-GHA and 6P4a-FANG systems indicate that more SRS sites were involved in the former. SRS3 and SRS4 motifs exhibited unique intermolecular interactions with deltamethrin and permethrin, respectively. Stronger binding affinity of 6P4a-GHA model was primarily attributed to interactions between deltamethrin and SRS-1, SRS-3, and SRS-6, as well as permethrin and SRS-1, SRS-4, and SRS-6.

A 250ns molecular mechanics thermodynamic integration simulation of the interaction between the CYP6P4a models and insecticides confirmed the docking binding affinity approximations, where 6P4a-GHA exhibited better affinity for deltamethrin (mmgb ΔG = −8.3 Kcal/mol) than 6P4a-FANG (mmgb ΔG = −8.15 Kcal/mol) (Table S3). Affinities for permethrin were also confirmed with 6P4a-GHA binding more strongly to permethrin (mmgb ΔG = −7.69 Kcal/mol) than 6P4a-FANG (mmgb ΔG = −7.56 Kcal/mol) (Table S3). While these differences in pyrethroid binding free energy between the wild-type and mutant alleles were modest, the strength of these data lies in the remarkable consistency between the thermodynamic integrations and molecular docking results.

### PLIF reveals possible mechanisms for preferential pyrethroid binding to 6P4a-GHA model

Analyses of protein-ligand interaction fingerprints (PLIF) allowed the identification of key substrate recognition site (SRS) residues and ligand pharmacophores responsible for the preferential binding of pyrethroids to the 6P4a-GHA mutant compared to the wild-type model (6P4a-FANG). Mutations in the 6P4a-GHA model (*M*^220^*I, V*^221^*F, V*^228^*M, L*^406^*F, V*^407^*I,* and *L*^409^*I*) induced changes in the active site architecture within SRS regions, leading to decreased efficacy of pyrethroid insecticides. The 6P4a-GHA system had more engaged SRS sites (SRS1–4 and SRS-6) than 6P4a-FANG, with SRS3 and SRS4 motifs forming interactions with deltamethrin and permethrin, respectively (Figs. 2E and 2F). Stronger binding affinity in the 6P4a-GHA model was predicted to be primarily induced by deltamethrin interactions with hotspot residues SRS-1 (*S*^119^*, H*^121^), SRS-3 (*T*^241^), and SRS-6 (*F*^495^), as well as permethrin bonding with SRS-1 (*R*^107^*, H*^121^), SRS-4 (*L*^314^), and SRS-6 (*F*^495^) (Figs. 2E and 2F). Deltamethrin’s bromo-ethenyl group forms a backbone hydrogen bond with *S*^119^, along with arene-H hydrophobic interactions between deltamethrin phenyl-ring and *H*^121^ of SRS-1 motif (Figs. 2E and 2F). For permethrin, the key interactions within SRS-1 predicted to influence the preference of the mutant 6P4a-GHA system are the arene attraction between *R*^107^ and permethrin’s phenyl ring, and *H*^121^ sidechain hydrogen bond donation to permethrin’s carboxylic-ester group (Figs. 2E and 2F). Structural changes induced by the 6P4a-GHA model mutations in the SRS-6 region, particularly at residue *F*^495^, enhanced substrate selectivity for permethrin. *F*^495^ formed arene-H interactions with methyl-ester and dimethyl-cyclopropane groups of permethrin, contributing to its binding affinity in the mutant enzyme. Furthermore, the 2-D interaction maps reveal that metabolism of deltamethrin by the mutant allele is driven by hydrogen bonding and arene-π electron interaction while binding to permethrin is mainly through arene-arene, arene-H, and arene-cationic hydrophobic interactions (Fig. S4). This was reflected in the binding affinities, where the mutant allele exhibited more affinity for deltamethrin than for permethrin (Table S3).

Docking ligands to Ghana *6P4b-^E^*^284^ (6P4b-GHA) and *6P4b-T*^291^*V*^294^*Y*^399^ (6P4b-FANG) produced inconsistent affinity and binding mode differences and was not further investigated.

### *CYP6P4a* gene is duplicated in Ghana

Recent studies have highlighted gene amplification of P450s as an insecticide resistance response mechanism. Here, Gene duplication of *CYP6P4a* was confirmed through copy number variation studies using qPCR (Fig. S5). A two-fold copy number of *CYP6P4a* was observed in the Ghanian *An. funestus* compared to the susceptible FANG laboratory strain which supplementing the previous report of an insertion between *CYP6P5* and *CYP6P4a* in Ghana. Quantification of copy number for *CYP6P4b* was challenging due to the non-specificity of the primers to amplify the Ghanian samples.

### Recombinant *An. funestus* CYP6P4a and CYP6P4b enzymes metabolise pyrethroids insecticides

We investigated the enzymatic activity of recombinant CYP6P4a and CYP6P4b enzymes in metabolising pyrethroid insecticides to validate their direct involvement in the phenotype and the impact of allelic variation on enzyme activity. Optimal expression of the enzymes was achieved 22 hours post-induction with IPTG (Fig. S6A). Substrate disappearance assays (Fig. S6B) revealed that the Ghanaian *6P4a-I*^220^*F*^221^*M*^228^*F*^406^*I*^407^*I*^409^ (6P4a-GHA) and Malawian *6P4a-L*^488^ (6P4a-MWI) significantly metabolised permethrin (P < 0.05 and P < 0.01, respectively), with 34.34% and 41.33% depletion respectively, compared to 6P4a-FANG with 27.57% depletion of the insecticide (Fig. 3A). For Type II pyrethroids, 6P4a-GHA and 6P4a-MWI demonstrated significantly higher depletion of deltamethrin, depleting 47.88% (P < 0.0001) and 69.59% (P < 0.0001), respectively, compared to 21.59% depletion by the 6P4a-FANG. This corresponds to 2.2-fold and 3.2-fold respectively, of elevated metabolic activity by the resistance alleles. Similarly, for alphacypermethrin, 6P4a-GHA and 6P4a-MWI significantly depleted the insecticide by 26.04% (3.7-fold) and 35.13% (4.9-fold) respectively, compared to the FANG variant that exhibited minimal metabolism (7.1%) (Fig. 3A). Similar patterns were observed for CYP6P4b enzymes, although to a lesser extent. For permethrin, the Ghanaian *6P4b-E*^284^ (6P4b-GHA), the Mozambican *6P4b-H*^26^ (6P4b-MOZ) and Ugandan *6P4b-N*^288^ (6P4b-UGA) exhibited depletions of 21.20%, 20.0%, and 19.11%, respectively, with no significant difference when compared to FANG *6P4b-T*^291^*V*^294^*Y*^399^ (6P4b-FANG) 16.22% depletion (Fig. 3B). With deltamethrin, 6P4b-GHA demonstrated a significant depletion of 30.05% (P = 0.001), 6P4b-MOZ exhibited 22.73% depletion (P = 0.0004), the 6P4b-UGA showed 13.30% depletion, while 6P4b-FANG exhibited little or no metabolism of the insecticide with only 3.15% depletion. This corresponds to an approximate 10-fold, 7.2-fold, and 4.2-fold elevated metabolic activity by the resistance enzymes over the FANG enzymes. The enzymes also metabolised alphacypermethrin, with 6P4b-GHA, 6P4b-MOZ, 6P4b-UGA and 6P4b-FANG producing 27.5%, 22.2%, 12.8%, and 17.7% depletion, respectively, with a significant difference obtained for the 6P4b-GHA vs. 6P4b-FANG comparison (P < 0.04) (Fig. 3B).

**Figure 3:**
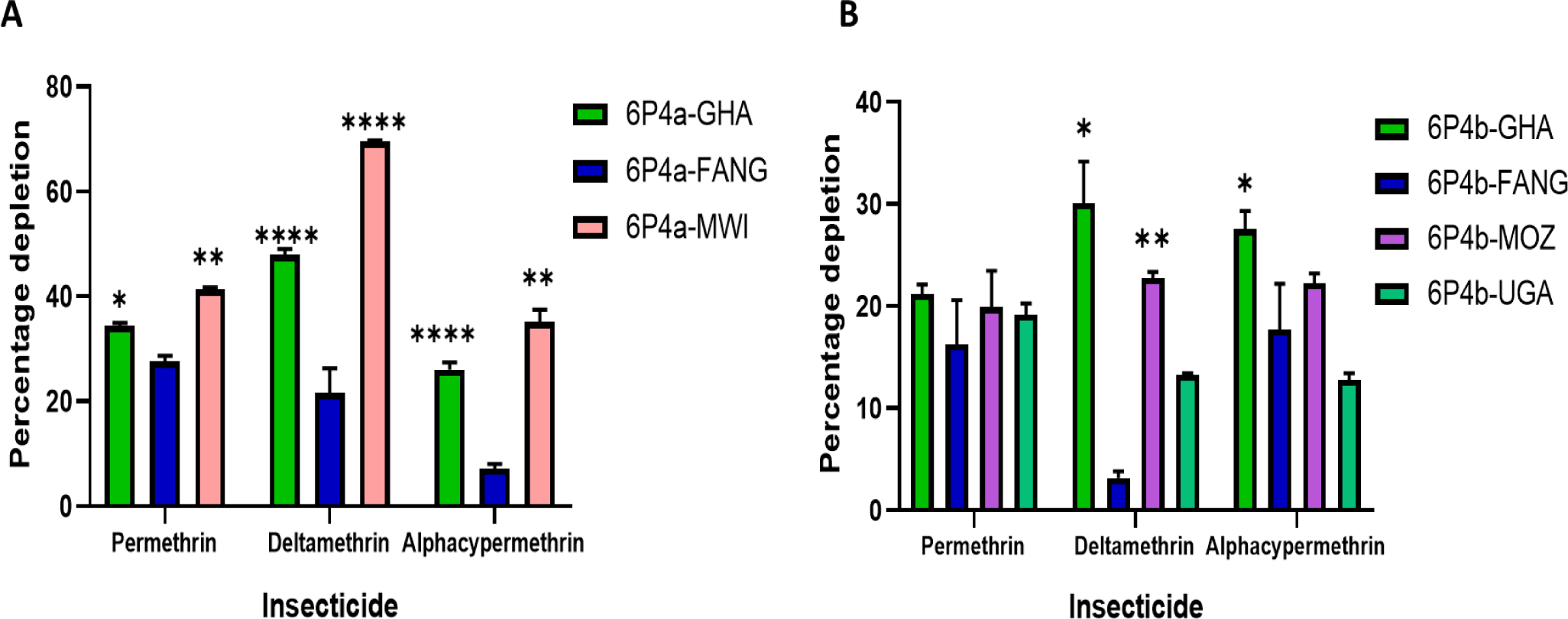
*In vitro* assessment of pyrethroid metabolism by CYP6P4a and CYP6P4b candidate alleles. **(A)**, CYP6P4a recombinant enzymes metabolism of pyrethroids; **(B),** CYP6P4b recombinant enzymes metabolism of pyrethroids. Values are mean ± SEM of three experimental replicates compared with negative control without NADPH. Depletion by recombinant resistant enzymes is significantly different from depletion by FANG recombinant enzymes at * p<0.05, ** p<0.01, *** p<0.001 and **** p<0.0001.

### *In vivo,* allelic variation and overexpression of *CYP6P4a* and *CYP6P4b* confer insecticide resistance

Given the overexpression of *CYP6P4a* and *CYP6P4b* in field populations, transgenic drosophila flies were constructed to assess the impact of the independent and sole over-expression of each of the genes on resistance to pyrethroids and further, to investigate the impact of allelic variation on resistance in an *in vivo* system. Quantitative RT-PCR analysis confirmed the significant overabundance of CYP6P4a and CYP6P4b transcripts in the transgenic *D. melanogaster* compared to non-transgenic control strains (Fig. S7). With 24h exposure to permethrin, GAL4-CYP6P4a-GHA flies and GAL4-CYP6P4a-MWI showed significantly lower mortality rates at different time intervals compared to the GAL4-CYP6P4a-FANG flies and the non-transgenic group (Fig. 4A). Also, averagely, a significant higher resistance was observed in the experimental group compared to the control group and GAL4-CYP6P4a-FANG flies throughout the 24h while revealing no difference in mean mortality between GAL4-CYP6P4a-FANG and the control group (Fig. 4G). This implies that at any given time of exposure, the experimental group was more resistant than the non-transgenic group and the overexpression of the GAL4-CYP6P4a-FANG allele confers no significant resistance to permethrin (Fig. 4G). Similarly, GAL4-CYP6P4a-GHA and GAL4-CYP6P4a-MWI flies were significantly more resistant to deltamethrin than GAL4-CYP6P4a-FANG and the control group at all time points (Fig. 4B). notably, after 12 h of deltamethrin exposure, GAL4-CYP6P4a-GHA and GAL4-CYP6P4a-MWI flies recorded 29% and 33.67% mortality, while GAL4-CYP6P4a-FANG and the control exhibited 61% and 70% mortality, respectively (Fig. 4B). Data also revealed a significantly high resistance in the experimental group compared to the control group and GAL4-CYP6P4a-FANG flies at all six different times, spanning 1h – 24h while no difference was recorded between the control group and the GAL4-CYP6P4a-FANG flies (Fig. 4G). For alphacypermethrin, the GAL4-CYP6P4a-GHA flies exhibited significant resistance to the insecticide compared to the GAL4-CYP6P4a-FANG and control groups, but overexpression of the MWI allele conferred significantly more resistance to alphacypermethrin compared to the control group only after 1h exposure (Fig. 4C). However, data showed a significantly high resistance in the GAL4-CYP6P4a-MWI compared to the control group when comparing mean mortalities for all six time points (36.59% vs. 45.66%) (Fig. 4G). Similarly, GAL4-CYP6P4b-GHA and GAL4-CYP6P4b-MOZ transgenic lines exhibited more resistance compared to GAL4-CYP6P4b-FANG and control flies. After 12 hours of permethrin exposure, GAL4-CYP6P4b-GHA and GAL4-CYP6P4b-MOZ flies exhibited significantly lower mortalities (36.05% and 30.73%, respectively), than the GAL4-CYP6P4b-FANG and control flies (44.35% and 59.17% respectively) (Fig. 4D). Mean mortalities for all six time points were 21.73%, 25.45%, 30.70%, and 41.38% for GAL4-CYP6P4b-GHA, GAL4-CYP6P4b-MOZ, GAL4-CYP6P4b-FANG flies, and the non-transgenic group, respectively (Fig. 4H). With deltamethrin, GAL4-CYP6P4b-GHA and GAL4-CYP6P4b-MOZ transgenic lines displayed lower mortality rates (35.56% and 28.30%, respectively) after 12 hours of exposure, while GAL4-CYP6P4b-FANG and the control recorded 51.93% and 70.74% mortality, respectively (Fig. 4E). For alphacypermethrin, GAL4-CYP6P4b-GHA and GAL4-CYP6P4b-MOZ exhibited higher resistance with mortalities of approximately 21.96% and 30.83%, respectively, compared to the control group with mortalities of 54.5% and 65.9% after 12 hours of exposure (Fig. 4F). Average mortalities for all six time points indicated that all transgenic groups were more resistant than the control group except for GAL4-CYP6P4b-FANG flies’ exposure to permethrin (Fig. 4H). These results collectively indicate that the upregulation of *CYP6P4a* and *CYP6P4b* confers resistance to pyrethroids but with allelic variation being a key factor in the phenotype as overexpression of the FANG variants did not confer resistance.

**Figure 4:**
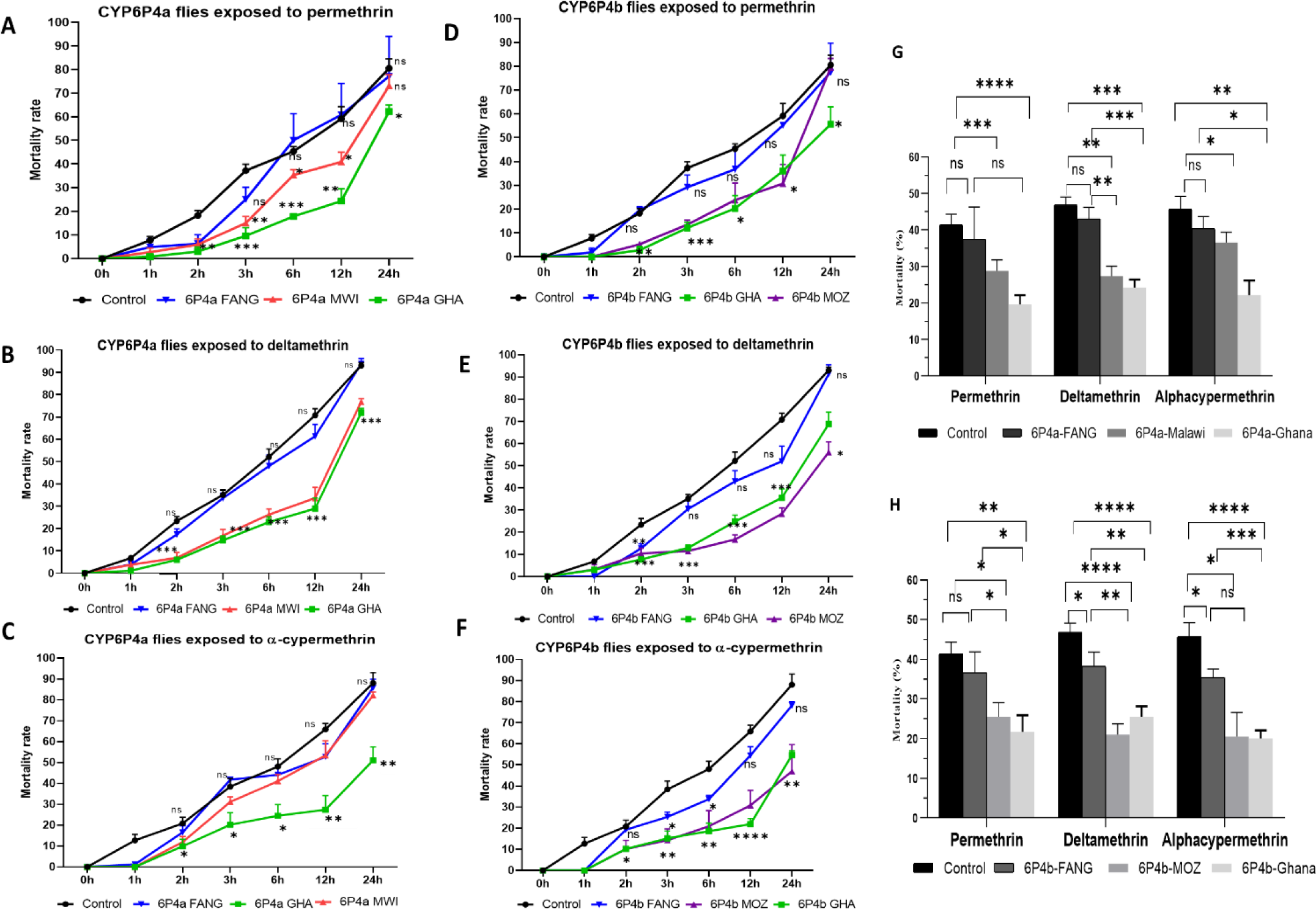
24h Insecticide contact bioassays with transgenic flies. over-expressing *CYP6P4a* alleles exposed to **(A),** permethrin; (**B),** deltamethrin; **(C),** alphacypermethrin, and transgenic flies over-expressing *CYP6P4b* alleles exposed to **(D),** permethrin; **(E),** deltamethrin; **(F),** alphacypermethrin. Data shown is mean ± S.E.M, significant difference in mortality is indicated for the GHA vs control and FANG vs control comparisons. Cumulative mortalities of transgenic flies and control flies to insecticides for all six time points **(G),** For CYP6P4a transgenic fly experiment; **(H),** For CYP6P4b transgenic fly experiment. * p<0.05, ** p<0.01, *** p<0.001 and **** p<0.0001.

### Two new molecular markers to track *CYP6P4a* and *CYP6P4b-* mediated pyrethroid resistance

Polymorphism analyses of full-length cDNA sequences allowed the identification of key mutations that were uniquely found in the Ghanaian population and absent in FANG. For *CYP6P4b,* the C to A nucleotide change at position 852 (Fig. S8A) led to amino acid change from aspartic acid (GA<u>C</u>) to glutamic acid (GA<u>A</u>). This mutation was used to design a DNA-based diagnostic tool (CYP6P4b-D284E) following the Amplification Refractory Mutation System (ARMS) PCR technique (Fig. 5A). For *CYP6P4a*, six mutations were identified: M220I, V221F, V228M, L406F, V407I, and L409I. Among these, the methionine-to-isoleucine change at position 220 (M220I) (Fig. S8B) was selected due to its proximity to substrate recognition site 2 (SRS-2) (Fig. S2A) and its role in the binding pocket of the enzyme, as revealed by molecular docking simulations. A Lock Nucleic Acid (LNA) probe-based PCR diagnostic tool (CYP6P4a-M220I) was designed targeting this mutation (Fig. 5B). The detailed protocols for the assay can be found in tables S4 and S5.

**Figure 5:**
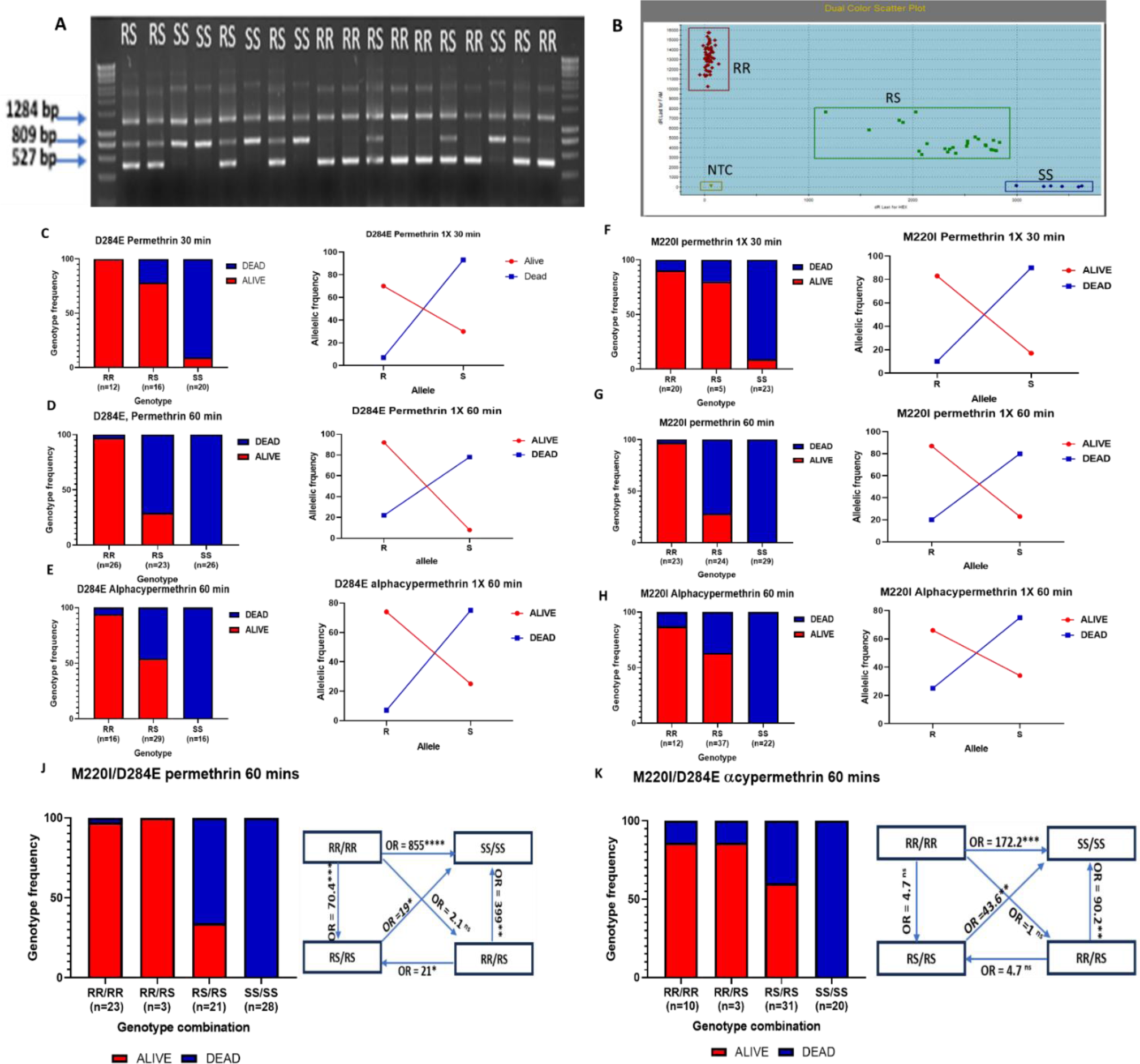
Design of DNA-based molecular diagnostic tools for detection of *CYP6P4a* and *CYP6P4b* resistance alleles and impact on resistance phenotype. (**A)** ARMS-PCR assay for genotyping the CYP6P4b-D284E marker; **(B), LNA** probe-based assay for genotyping the CYP6P4a-M220I marker. Association of the CYP6P4b-D284E and the CYP6P4a-M220I mutations with insecticide resistance phenotype. Distribution of CYP6P4b-D284E resistance marker among F3 FANGxGHANA hybrid *An. funestus* mosquitoes exposed to **(C),** Permethrin for 30 mins; **(D),** Permethrin for 60 mins**; (E),** Alphacypermethrin for 60 mins. Distribution of CYP6P4a-M220I resistance marker among F3 FANGxGHANA hybrid *An. funestus* mosquitoes exposed to **(F),** Permethrin for 30 mins, **(G),** Permethrin for 60 mins; **(H),** Alphacypermethrin for 60 mins. 284E, 220I (R) and D284, M220 (S) allele frequency distributions between Alive and Dead mosquitoes are shown in line plots. Combined impact of CYP6P4b-D284E and CYP6P4a-M220I mutations on pyrethroid insecticide resistance. Distribution of genotypes in FANGxGHANA hybrid An. funestus mosquitoes exposed to (J), permethrin for 60 mins; (K), alphacypermethrin for 60 mins.

### *CYP6P4a* and *CYP6P4b* markers are strongly associated with pyrethroid insecticide resistance

Assessing the genotypic composition of the field mosquitoes in Ghana revealed a high prevalence of the markers in the population. Genotyping the CYP6P4a-M220I marker revealed that 69.5% of individuals (32/46) were homozygous 220I/220I (RR), 21.74% (10/46) were heterozygous M220/220I (RS), and 8.7% (4/46) were homozygous susceptible M220/M220 (SS) (Fig. S8C). This corresponds to an allelic frequency distribution of 79.2% and 20.8% for the R and S alleles in the population. For the CYP6P4b-D284E marker, a higher percentage of the resistance allele was recorded; 9% (44/46) 284E/284E (RR) and 4.35% (2/46) D284/284E (RS) genotypes, with no D284/D284 (SS) individual. This corresponds to a 97.8% frequency of the R allele and 2.2% frequency of the S allele in the population (Fig. S8C). In contrast, all FANG samples were homozygous SS for both markers. The high pyrethroid resistance and high frequency of the resistant alleles in the field population impeded the establishment of a genotype-phenotype correlation. We opted for a genetic cross strain between the highly resistant Ghanaian mosquitoes and the fully susceptible laboratory strain FANG to establish a population of mosquitoes with all three genotypes to allow genotype/phenotype correlation studies. At the F3 generation, the population was made of 25% RR, 35% RS, and 40% SS individuals for the CYP6P4b-D284E marker (Fig S8D). This confirmed that the genetic cross was successfully segregated the genotypes. These mosquitoes were exposed to pyrethroids using the WHO insecticide bioassays for the establishment of the genotype-phenotype association. Exposing mosquitoes to permethrin 1X for 30 mins resulted in 61% mortality, while after 1 hour, mortality was 85 ± 2.57% for permethrin 1X and 54 ± 2.57% for alphacypermethrin 1X (Fig. S8E). These mortalities are far higher than the 11.6±5% for permethrin 1× and 1.25±1.25% for alpha-cypermethrin 1x, previously recorded in the field mosquitoes in the same year ^21^, highlighting the potential association between these markers and the phenotype.

#### CYP6P4b-D284E

86 % of highly susceptible (HS) mosquitoes (dead after 30-minute exposure to permethrin) were homozygous susceptible SS (18/21) and 14 % heterozygotes (3/21) with no homozygous resistant RR individuals observed (Fig. 5C left). In contrast, highly resistant mosquitoes (HR) that survived 60-minute exposure to permethrin were predominantly homozygous RR (25/30), and heterozygotes (5/30), constituting 83% and 17% of the HR mosquitoes respectively, with no SS individual (Fig. 5D left). Allele frequency distribution further supported the trend where in HR mosquitoes, 97% R and 20% S allele frequencies were recorded, while in the HS mosquitoes, the R allele was present at 7% and the S allele was present at 93% (Fig 5C right and 5D right). A strong positive association was established between CYP6P4b-D284E marker and ability to survive exposure to permethrin, with RR individuals having greater chance of surviving compared to SS individuals (OR = 901; CI = 35.06 to 23156.34; P < 0.0001, Fisher’s exact test). (Table S6). Similar observations were made for mosquitoes exposed to alphacypermethrin for 60 mins, where carrying the RR genotype enhanced alphacypermethrin resistance, reflected in the significant odds ratios observed in the RR vs. SS comparison (OR = 341; CI = 12.09 to 9016.19; P < 0.0001, Fisher’s exact test) and RS vs. SS comparison (OR = 40.33; CI = 2.21 to 735.99; P = 0.0126, Fisher’s exact test) (Fig 5E left). These findings were further supported by the allele frequency distribution, with the R allele occurring at 74.2% in surviving mosquitoes and 25.8% in the dead mosquitoes, while the S allele frequency was 25% in the surviving mosquitoes and 75% in the dead mosquitoes (Fig.5E right). Overall, for resistance to both type 1 and 2 pyrethroids, an additive effect was observed as RR individuals survived significantly more than RS individuals (Table S6).

#### CYP6P4a-M220I

88% of the highly susceptible mosquitoes (HS) (dead after 30-minute exposure to permethrin) were SS (21/24), 4% RS (1/24) and 8% RR (2/24) (Fig. 5F left). In contrast, the mosquitoes that survived after 60 minutes exposure to permethrin were predominantly RR (22/30) and RS (8/30), with no SS individual (Fig. 5G left). Furthermore, the R allele was more prevalent among the survivors (87%), while the S allele was more common among the dead mosquitoes (80%) (Fig. 5G right). These findings established a strong association between the CYP6P4a-M220I marker and permethrin resistance with highly significant odds ratio when comparing RR vs. SS (OR = 885; CI = 34.41 to 22763.86; P < 0.0001), Fisher’s exact test) and RS vs. SS (OR = 30.4; CI = 1.65 to 560.83; P = 0.0217, Fisher’s exact test). Similar results were obtained with alphacypermethrin as alive mosquitoes were predominantly RR (10/31) and RS (21/31), with no SS individual (0/30) while the dead were predominantly SS (22/40) and RS (16/40), with 2 RR individuals (2/40). Significant odds ratios were observed for the RR vs. SS (OR = 189; CI = 8.32 to 4295.30; P = 0.001, Fisher’s exact test) and RS vs. SS (OR = 58.64; CI = 3.81 to 1039.32; P = 0.0055, Fisher’s exact test) comparisons (Fig. 5H left). This was supported by allele frequency distributions, with the R allele occurring at 66% in the alive mosquitoes and 75% S allele in the dead (Fig. 5H right).

#### CYP6P4a-220I and CYP6P4b-284E combined to confer higher resistance levels

Double homozygote resistant genotype (RR/RR) conferred significantly more resistance to permethrin while double heterozygous genotype (RS/RS) conferred resistance but to a lower extent. For type II pyrethroid (alphacypermethrin), RR/RR and RS/RS genotypes conferred similar resistance levels (Table S6). Specifically, individuals that survived permethrin exposure were more of the RR/RR (22/30) accounting for 73.3%, RR/RS (3/30) representing 10% and RS/RS (5/30) representing 16.7% of the alive mosquitoes. The majority of individuals dead after exposure were SS/SS (28/45) making up 62.2%, and RS/RS (16/45) with only one RR/RR individual (2.2%) (Fig. 5J). A strong genotype phenotype correlation was therefore established in the RR/RR vs SS/SS comparison (OR = 855; P < 0.0001, Fisher’s exact test) as well as the other genotype combinations (Table S6). The additive effect of the markers was also observed with RR/RR individuals surviving significantly more than RS/RS individuals (OR = 70.4; CI = 7.48 to 662.33; P = 0.0002) (Fig. 5J right, Table S6). For alphacypermethrin, all alive individuals had at least one resistant allele, while the dead were more of the double homozygous susceptible genotype (Fig. 5K). Having both CYP6P4a-220I and CYP6P4b-284E resistance alleles in the RR/RR and RS/RS states significantly increased resistance to alphacypermethrin compared to the double homozygote susceptible (SS/SS) state (OR = 172.2; CI = 7.56 to 3923.94; P = 0.0012 and OR = 43.6; CI = 2.43 to 785.21; P = 0.0104, respectively). The analysis of resistance to alphacypermethrin revealed that the impact of RR/RR genotype on resistance was not significantly different from other genotype combinations. This suggests that individuals do not necessarily require the RR/RR genotype to exhibit high resistance to alphacypermethrin (Fig. 5K right, Table S6).

### *CYP6P4a* and *CYP6P4b* resistance alleles reduce the efficacy of pyrethroid-based tools

Cone assays performed with different LLINs using F4 FANG/GHANA mosquitoes revealed mortalities rate of 30.45 ± 4.94%, 44.09 ± 6.57%, and 32.44 ± 5.74% respectively with PermaNet 2.0, DuraNet and Olyset (Fig. S8F).

#### CYP6P4b-D284E

Assessing the correlation between genotypes and survivorship revealed that RR mosquitoes were significantly more able to survive exposure to PermaNet 2.0 net (Deltamethrin-based) than SS mosquitoes (OR=5.8; CI=1.49 to 22.01; P<0.0001) (Fig 6A). A similar observation was made for RS mosquitoes which survived more than SS (OR=5.4; CI=1.6579 to 17.6481; P = 0.0051). Although mosquitoes that survive had a higher frequency of RR than RS genotype, the difference was not significant (OR=2.6; CI=0.6034 to 10.9184; P=0.202) (Table S5) confirming the pattern observed for WHO tube test with alphacypermethrin where bearin. The impact was even greater on Olyset (permethrin) efficacy (Fig 6B) where homozygous RR and heterozygous RS individuals significantly survived better than homozygous SS individuals (OR 35.8; CI = 6.33 to 202.37; P<0.0001 and OR 14.0; CI = 3.40 to 57.11; P = 0.0003). Analogous observations were made for DuraNet with RR and RS individuals exhibiting more resistance than SS individuals (Table S5, Fig 6C).

**Figure 6:**
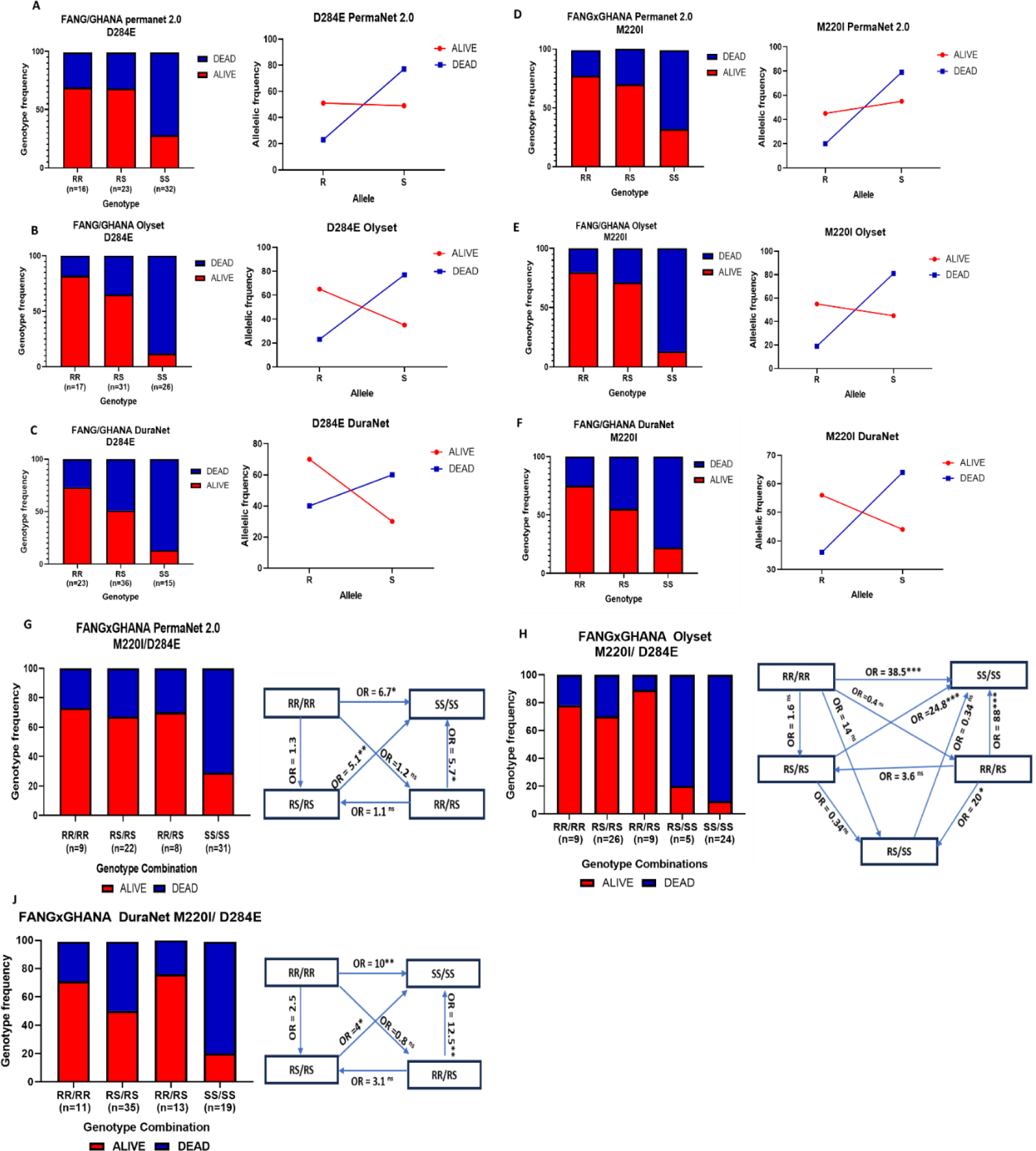
Genotype/ phenotype Association of the CYP6P4b-D284E and the CYP6P4a-M220I mutations with pyrethroid bed net efficacy. Distribution of CYP6P4b-D284E resistance mutation genotypes among resistant and susceptible F4 FANGxGHANA hybrid *An. funestus* mosquitoes exposed to: **(A),** PermaNet 2.0 for 3 mins; **(B),** Olyset for 3 mins; **(C),** Duranet for 3 mins. Distribution of CYP6P4a-M220I resistance mutation genotypes among resistant and susceptible F4 FANGxGHANA hybrid *An. funestus* mosquitoes exposed to: **(D),** PermaNet 2.0 for 3 mins; **(E),** Olyset for 3 mins; **(F),** DuraNet for 3 mins. Combined impact of CYP6P4b-D284E and CYP6P4a-M220I mutations on pyrethroid bednet efficacy. Distribution of genotypes in FANGxGHANA hybrid *An. funestus* mosquitoes exposed to: **(G),** PermaNet 2.0 for 3 mins; **(H),** Olyset for 3 mins; **(J),** DuraNet for 3 mins. R and S Allele frequency distributions between Alive and Dead mosquitoes after exposure are shown in line plots.

#### CYP6P4a-M220I

Genotyping alive and dead mosquitoes after exposure to PermaNet 2.0 also revealed that CYP6P4a-220I homozygous resistant mosquitoes (RR) were significantly more able to survive exposure to PermaNet 2.0 nets than the homozygous susceptible CYP6P4a-M220 mosquitoes (SS) (OR = 6.7; CI = 1.36 to 34.35; P = 0.0205). Similarly, heterozygote mosquitoes survived better than susceptible SS (OR=4.67; CI=1.66 to 14.50; P = 0.0054) (Fig 6D). Also, RR and RS genotypes significantly increased chances of surviving exposure to Olyset compared to SS genotype (OR = 27; P < 0.0001; and OR = 16.57; P < 0.0001) (Fig. 6E). Similarly, double homozygous RR and heterozygous mosquitoes survived more after coming in contact with DuraNet than the homozygous SS mosquitoes (OR = 10.5; P = 0.01 and OR = 4.295; P <0.05, respectively) (Fig. 6F).

#### Combined effect of both markers on pyrethroid bednet efficacy

An independent segregation of genotypes at both genes was observed at F4, with several genotype combinations obtained, including: RR/RR, RR/RS, RS/RS, RS/SS and SS/SS. Genotype/phenotype comparisons revealed that RR/RR, and RS/RS mosquitoes survived exposure to PermaNet 2.0 significantly more than SS/SS (OR = 6.7; CI = 1.18 to 37.79; P = 0.032 and OR = 5.1; CI = 1.55 to 16.71; P = 0.0073, respectively) (Fig. 6G). With Olyset, RR/RR, RR/RS, and RS/RS individuals had far greater ability to survive relative to SS/SS individuals (OR 38.5; P = 0.0008, OR=24.8; P=0.0002, OR=88; P=0.0005, respectively) and (Fig. 6H, Table S5). A similar pattern was observed for exposure to DuraNet, where RR/RR, RR/RS and RS/RS individuals also survived significantly more than SS/SS individuals (OR = 10; CI = 1.78 to 56.15; P = 0.009; OR = 12.5; CI = 2.29 to 68.25; P = 0.0035, OR = 4; CI = 1.10 to 14.38; P = 0.036, respectively) (Fig. 6J).

### Africa-wide distribution of *CYP6P4a* and *CYP6P4b* resistance alleles

Screening countries across different regions of Africa, including West, East, Central, and South Africa for the presence of the CYP6P4b-D284E and CYP6P4a-M220I resistance markers, partially established the geographical spread of the markers on the continent. The CYP6P4b-284E resistance marker was close to fixation in Ghana (98%) and Sierra Leone (91%), at an intermediate frequency in Benin (51%), all West African countries, but absent in other countries including Cameroon and the DRC in Central Africa, Uganda and Tanzania in East Africa, and Mozambique and Malawi in Southern Africa (Fig. 7A). Also, the CYP6P4a-220I resistance marker was also identified in Ghana and Seirra Leone at high frequency (80% and 91%, respectively) and afore-mentioned countries. (Fig. 7B). Assessment of the evolution of the markers between 2014, 2021, and 2022 in the sampled Ghana population revealed that the markers have increased in the population between the two time points (79-86-98% for CYP6P4b-284E and 75-80% for CYP6P4a-220I), with high chances of getting fixed in the population in a few generations (Fig. 7C and 7D).

**Figure 7.**
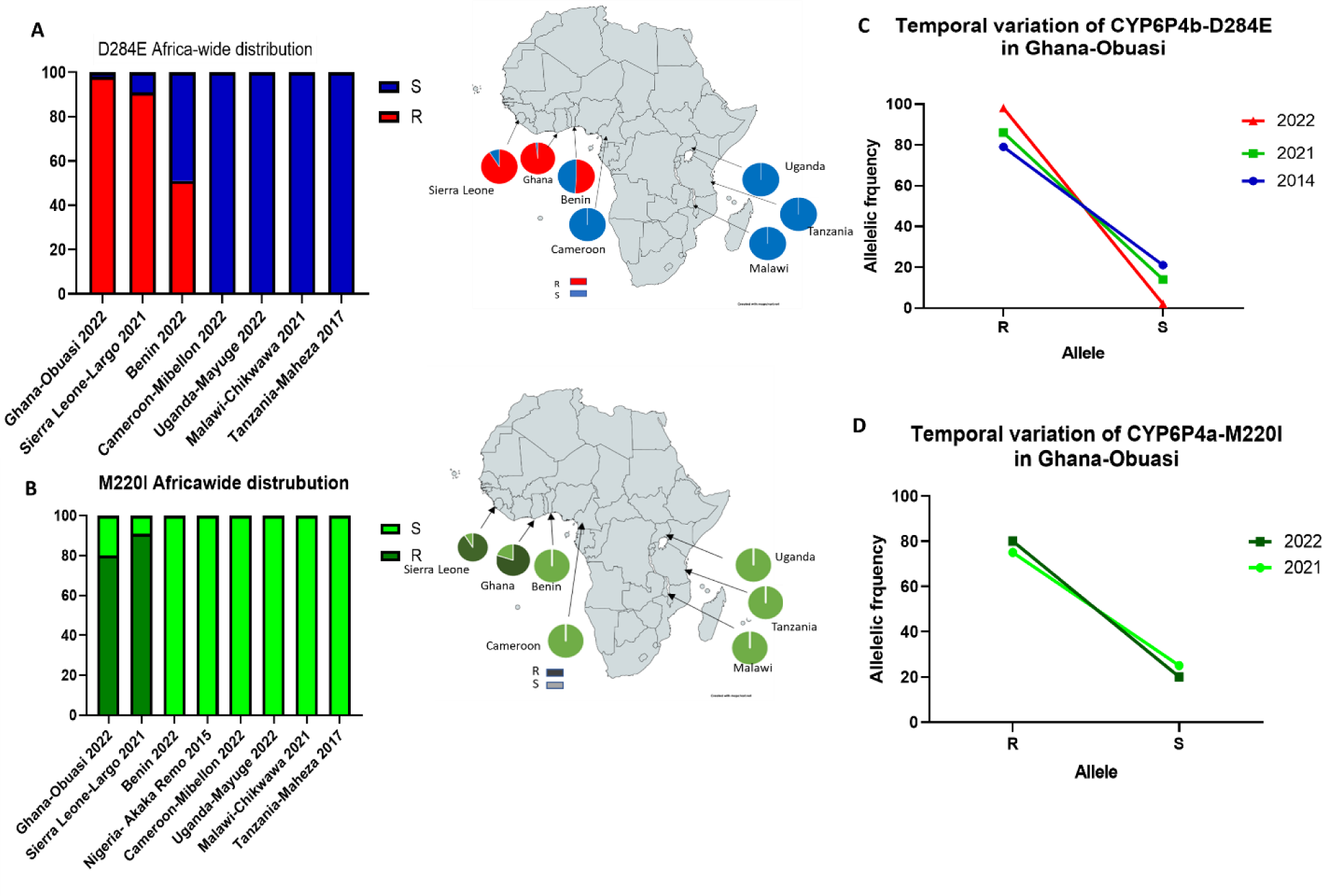
Geographical distribution of molecular markers across Africa. **(A),** CYP6P4b-D284E resistance allele **(B),** CYP6P4a-M220I resistance allele across Africa. **(C),** Temporal assessment of the allele frequency distribution of the CYP6P4b-D284E resistance marker in the *An. funestus* population of Ghana-Obuasi. **(D),** Temporal assessment of the allele frequency distribution of the CYP6P4a-M220I resistance marker in the *An. funestus* population of Ghana-Obuasi.

## Discussion

In this study, we elucidated the genetic basis of metabolic resistance to pyrethroids in west African populations of the major malaria vector *An. funestus* establishing that *CYP6P4a* and *CYP6P4b* are driving pyrethroid resistance through overexpression and allelic variation. Furthermore, we introduce the two field-applicable DNA-based diagnostic markers for detecting P450-based metabolic resistance to pyrethroids in *An. funestus* populations in West Africa to facilitate the detection, monitoring and management of insecticide resistance.

### Directional selection of *CYP6P4a* and *CYP6P4b* is associated with pyrethroid resistance

Analysing the sequences of representative populations of the four regions of Africa revealed an overall reduced diversity of *CYP6P4a* and *CYP6P4b*. The sampled populations confirmed the presence of four major haplotypes, forming four geographical clusters: southern Africa (Mozambique/Malawi), in East-Central Africa (Uganda/Cameroon), Benin (West), and Ghana alone (West). This population clustering is interestingly very close to what was observed in the study of Mugenzi et al ^14^ where sequencing of promoter regions of *CYP6P9a* revealed the existence of haplotypes forming four geographical clusters: southern Africa, East-Central Africa (Kenya/Uganda/Cameroon), West-Central Africa (Benin/DRC), and in Ghana only (West Africa). This further confirms that pyrethroid resistance in *An. funestus* is driven by the selection and overexpression of the genes found in the *rp1* quantitative trait locus (QTL), a locus that has been shown to account for more than 85% of pyrethroid resistance in *An. funestus* ^22,23^. In Ghana, a notable increase in the frequencies of the *6P4a-I*^220^*F*^221^*M*^228^*F*^406^*I*^407^*I*^409^ major allele from 7% to 78% and the *6P4b-E*^284^ major allele from 44% to 100*%* for samples collected in the years 2014 and 2021 was observed. This increase coincides with the significant decrease in pyrethroid susceptibility from 36.11 ± 3.87% mortality in 2014 ^24^ to 11.6 ± 5% in 2021 ^21^, for the same locality (Atatam, Ghana). The directional selection of these genes under insecticide pressure is similar to the reduced diversity observed in genes found in the *rp1* locus like the tandemly duplicated *CYP6P9a* and *CYP6P9b* (22), but contrasts with the high diversity of genes not found in the *rp1* like the *CYP325A* gene involved in pyrethroid resistance in Cameroon ^26^ and *CYP6M7*, driving pyrethroid resistance in Zambia ^27^, both found in the *rp2* QTL, and *CYP9J11* found in the *rp3* QTL^28^. In the southern African population, a remarkable low diversity was observed in 2014 with the existence of major alleles already, which became fixed in 2021. However, the lower expression of the genes relative to *CYP6P9a/b* ^29^ suggests that the alleles are not the major drivers of resistance and could have been selected along with the *CYP6P9a* and *CYP6P9b* genes-the major drivers of pyrethroid resistance in this region ^15,22^ via genetic hitchhiking. For Eastern (Uganda) and Central Africa (Cameroon), *CYP6P4a* and *CYP6P4b* were highly diversified, with no signs of selection in 2014. In 2021, there is the existence of a common major allele for each gene (*6P4a-R*^169^*E*^295^*K*^297^ and *6P4b-N*^288^) which could be as a result of the recent gene flow from Eastern to Central Africa^30^. The identification of directional selection and polymorphism in the *CYP6P4a* and *CYP6P4b* genes highlights the adaptive nature of mosquito populations in response to insecticide pressure. Continued monitoring of these genes and their variations will contribute to decipher mosquito evolution, gene flow, and the spread of resistance. This knowledge can inform future strategies for controlling and managing resistance in mosquitoes.

### The directionally selected *CYP6P4a* and *CYP6P4b* alleles mediate insecticide resistance via mechanisms involving gene duplication, allelic variation, and gene overexpression

#### The *CYP6P4a* is duplicated in Ghana

The confirmation of the duplication of *CYP6P4a* by qRT-PCR partially explains the overexpression of the gene and further enlightens on the composition of the previously reported 6.2kb insertion in Ghana ^31^. It could also account for the nearly 2X expression of *CYP6P4a* over *CYP6P4b* (44.8- and 23.9-fold) from published transcriptomic data ^15^. Interestingly, the duplication is peculiar to Western Africa (Ghana), as qPCR evaluation of gene duplication in Southern Africa had revealed no duplication, neither for *CYP6P4a* and *CYP6P4b* nor *CYP6P9a and CYP6P9b* in Malawian and Mozambican *An. Funestus*^29^. This 6.2kb insertion coming with the duplication of *CYP6P4a* could therefore be a response mechanism to insecticide pressure, and could contribute to the high pyrethroid resistance now observed in West African *An. funestus* population. Further study on the entire composition of the insertion would help evaluate its impact on the high pyrethroid resistance levels observed in the field. Although, overexpression of cytochrome P450 in mosquitoes is more often associated with changes in the cis- or trans-acting regulatory loci ^14,15,18,30^, the duplication of *CYP6P4a* here adds to growing evidence that copy number variation is also a major contributor. This has been shown recently in the other major malaria vector *An. gambiae* ^32,33^.

### Allelic variation enhances CYP6P4a and CYP6P4b pyrethroid detoxification activity

#### The *CYP6P4a* mutant allele has greater binding affinity for pyrethroids

*In silico* modelling and docking has been an essential tool in the characterisation of enzyme-substrate interactions of many P450s ^34–36^. The hydroxylation or hydrolysis of the 4’-phenoxybenzyl moiety along with the cis/trans methyl spot of type I (permethrin) ^37^ and type II (deltamethrin) ^25,38^ pyrethroids are reported to be the main routes for pyrethroid metabolism. In the implemented docking protocol of this study, the 4′ hydroxylation of the phenoxybenzyl moiety, which is the major pyrethroid metabolism route was indeed more enriched compared to other minor metabolic routes aligning with previously reported structural experiments ^38^. Molecular docking and thermodynamic integration simulation showed that the major *CYP6P4a* allele in Ghana (mutant) exhibits higher affinity for permethrin and deltamethrin than the susceptible lab strain FANG allele (wildtype), underscoring the key role that non-synonymous mutations on insecticide resistance. Mutations in the 6P4a-GHA model were predicted to induce structural backbone deviations in the active site architecture within SRS-1 and 6 regions, resulting in stronger binding affinity in the 6P4a-GHA model. SRS motifs in P450 enzymes play a crucial role in the metabolism of insecticides. Deltamethrin preference for hotspot residues SRS-1 (*S*^119^*, H*^121^) and SRS-6 (*F*^495^), as well as permethrin bonding with SRS-1 (*R*^107^, *H*^121^) and SRS-6 (*F*^495^) on the mutant (6P4a-GHA) allele were the main discriminative patterns in pyrethroids binding modes to the wild-type vs. mutant-type models and may suggest a causal link with the 6P4a-GHA mutations. In agreement with this, previous studies ^39^ investigated the significance of conserved SRS-1 CYP3A4 residues through site-directed mutagenesis. The findings highlighted the critical role of the highly conserved residue *S*^119^ in determining specificity in the active site topology in steroid 6beta-hydroxylation. Interestingly, both pyrethroids were predicted to bind more mutant SRS-1 residues than the wild-type. Also, the SRS-6 region of CYP6AE11-20 has been studied using *in vitro* metabolism and modelling techniques. It was found that amino acid substitutions at position 495 in SRS-6 led to significant alterations in the shape and chemical environment of P450 enzyme active site ^40^. Interestingly, our findings reveal that the structural changes induced by the 6P4a-GHA model mutations in the SRS-6 region, particularly at residue *F*^495^, result in a favourable conformation that enhances substrate selectivity for permethrin over the wild-type case. Similar substrate affinity was observed with the *An arabiensis CYP6P4* orthologue ^41^ but contrary to what was reported for the *Ae. albopictus* orthologue *CYP6P12* ^42^ where the 4′ hydroxylation of the phenoxybenzyl moiety was not a detoxification route with molecules adopting unfavourable poses. While most of these mutations mapped to locations within or adjacent to the active site, the CYP6P4b-284E mapped to an external loop at considerable distance away from the catalytic site, making it difficult to predict the functional impact of the mutation through molecular docking alone. However, mutations in surface residues have been shown to influence electron transfer rates from P450 reductase and/or cytochrome b5 ^43,44^, and this could be a potential mechanism involved with this mutation.

#### *In vitro, CYP6P4a* and *CYP6P4b* mutant alleles were better metabolisers of pyrethroids

The patterns of differential enzyme-substrate interactions revealed via molecular docking were confirmed with *in vitro* insecticide metabolism assay. Recombinant enzymes exhibited direct metabolism of types I and II pyrethroid, with more metabolism observed for deltamethrin as predicted by the molecular docking experiment. Metabolism chromatograms displayed distinct metabolite peaks, which we predict to correspond to deltamethric acid, cyano(3-hydroxyphenyl)methyl deltamethrate, and 4’-hydroxy-deltamethrin (Fig3A). These findings support the fact that the P450 enzymes metabolise pyrethroids preferentially via ring hydroxylation^38^. Nevertheless, the precise identification of the metabolites from *CYP6P4a/b* insecticide depletion requires further experimental procedures like mass spectroscopy and nuclear magnetic resonance. The Ghana *6P4a-I*^220^*F*^221^*M*^228^*F*^406^*I*^407^*I*^409^ and *6P4-E*^284^ alleles exhibited significantly higher depletion compared to the FANG variants *6P4a-N*^286^*R*^289^*S*^291^ and *6P4b-T*^291^*V*^294^*Y*^399^ the selection of this allele in the population may be associated to their increased metabolism of insecticides, imparting resistance. Gene orthologues have shown similar profiles as reported for the *CYP6P4* orthologue in *An. gambiae s.s.* metabolised type I and type II pyrethroids ^32^, whereas the *An. arabiensis CYP6P4* orthologue was found metabolise permethrin but not deltamethrin ^41^. Although the *CYP6P4a* Malawi allele exhibited higher activity, its contribution to resistance might be limited due to its lower expression level compared to *CYP6P9a*/b genes-the major drivers of pyrethroid resistance in southern Africa. Allelic variation as a mechanism in insecticide resistance has also been reported in *CYP6a2*, in *D. melanogaster* ^45^, *GSTe2* in association with DDT resistance in *An. funestus* ^13^ and *Ae. aegypti* ^46^, and *CYP6P9a*/*b,* in association with pyrethroid resistance in *An. funestus* ^25^. For *CYP6P4a*, the *I*^220^*F*^221^*M*^228^*F*^406^*I*^407^*I*^409^ haplotype found in Ghana accounts for the increased detoxification activity observed. Mutagenesis experiments could help identify the mutation(s) that account for the greatest difference in enzyme activity. For *CYP6P4b*, the *284E* mutation impacts enzyme activity and the simple reversal of the amino acid to D284 could help better quantify the biochemical impact of the mutation. Such experimental procedures were used to evaluate the impact of key mutations in *An. funestus CYP6P9a* and *CYP6P9b* ^25,47^ and in human *CYP2D6* ^48^.

#### *In vivo*, allelic variation and overexpression of *CYP6P4a* and *CYP6P4b* impact pyrethroid insecticide resistance

In West Africa, where *CYP6P4a* and *CYP6P4b* are overexpressed, the consequential impact of the overexpression of the mutant alleles in the field was validated, with higher resistance to pyrethroids recorded in the transgenic flies compared to the non-transgenics. The phenotypic impact of mutations under selection in this population was revealed as flies overexpressing the mutant alleles (Ghana) showed lower mortality compared to those overexpressing the wild-type (FANG). This shows that the selection of the allele was not random but rather due to its higher metabolic efficiency. These mechanisms are similar to those reported for the *CYP6P9a* and *CYP6P9b* sister genes ^25^ and also for *GSTe2* conferring DDT resistance ^13^, but in the *CYP6P12 Ae. albopictus* orthologue, it was demonstrated that overexpression of the gene conferred resistance to deltamethrin but increased mortality to permethrin ^42^. This study therefore validates that *CYP6P4a* and *CYP6P4b* together aggravate pyrethroid resistance in Ghana via mechanisms of allelic variation and overexpression that includes gene duplication.

#### The *CYP6P4b*-*D284E* and the *CYP6P4a*-*M220I* combine to exacerbate resistance to pyrethroids

The *CYP6P4b-D284E* and *CYP6P4a-M220I* molecular markers individually and in synergy, were strongly associated with pyrethroid resistance and a significant decrease in the effectiveness of pyrethroid-only nets (Olyset, PermaNet 2.0, and DuraNet), where homozygous mutants and heterozygotes for the two alleles survived more compared to wild-types. This is similar to the impact of *CYP6P9a* and *CYP6P9b* markers on the efficacy of pyrethroid bed nets ^14,15,18^. It is now imperative to assess the influence of these markers on the effectiveness of nets containing the synergist piperonyl butoxide (PBO) and on dual-action nets that combine pyrethroids with other insecticides in semi-field conditions, such as experimental huts to further evaluate the effectiveness of various control tools. Also, it is important to investigate the roles of these alleles in cross-resistance to other insecticide classes including organochlorines, organophosphates, carbamates, neonicotinoids, and insect growth regulator. In effect, studies on resistance to DDT in West Africa showed that CYP6P4a/b were the most overexpressed detoxification genes in DDT-resistant mosquitoes ^49^ as well as in carbamate-resistant mosquitoes^50^ but their implication in the phenotype needs to be validated. These evaluations will guide the selection of optimal insecticide-based vector control tools in regions where resistance is driven by *CYP6P4a/b*.

#### The *CYP6P4b*-*D284E* and the *CYP6P4a*-*M220I* alleles are prominent in West Africa

The new DNA-based markers designed here can now be effectively used in resistance management through routine molecular monitoring and tracking of resistance spread in the field. Analysis of the geographical distribution of these markers has revealed their high prevalence in sampled populations from Ghana, Sierra Leone, and Benin, with their prevalence steadily increasing over time. These mutations, however, remain absent in other regions of Africa, highlighting their association with West Africa and the restriction to gene flow among populations of this species between the major geographical regions ^16,31^. Indeed, the L119F-GSTe2 allele was found to be confined mainly to West Africa, low in Central and absent in Southern ^13^ whereas the CYP6P9a/- b resistant alleles were nearly fixed in Southern Africa but absent in other regions ^14,15,18^. It is now imperative to continue assessing the distribution of these markers in other West African countries where pyrethroid resistance is prevalent, while also maintaining continuous monitoring in other African regions for potential gene flow or de novo occurrence. This proactive approach will enable effective resistance management before the alleles reach high frequencies, thereby maintaining the efficacy of pyrethroid-based control tools and, ultimately, malaria elimination efforts.

## Conclusion

This study made use of a combination of computational analysis and laboratory experiments to provide valuable molecular diagnostic tools and in-depth insights into the molecular mechanisms by which the duplicated *CYP6P4a* and *CYP6P4b,* the most overexpressed detoxification genes in pyrethroid-resistant *An. funestus* in West Africa, drive pyrethroid metabolic resistance. The identification of markers within these genes that compromise the effectiveness of pyrethroid insecticide-based vector control tools represents a significant advancement in malaria vector control efforts in West Africa. Consequently, we encourage the deployment of nets incorporating piperonyl butoxide (PBO) or chlorfenapyr for enhanced impact, while also encouraging the development of alternative mosquito management strategies. Additionally, we emphasize the importance of conducting further investigations into the underlying mechanisms and genes associated with resistance. This knowledge will be instrumental in the development of improved insecticides that will enhance vector control measures.

## Methods

### Mosquito samples

For this study, mosquitoes were collected between 2018 and 2021 from the following localities: Mibellon in Cameroon (6° 4′ 60″N,11° 70′0″E)^55^; Mayuge in Uganda (0° 23′ 10.8″N, 33° 37′ 16.5″E)^56^; Obuasi in Ghana (06° 17.377″ N, 001° 27.545″ W) ^21^; Chikwawa in Malawi (16° 1′ S; 34° 47′ E) and Palmeira in Mozambique (25° 15′ 19′′ S; 32° 52′ 22′′ E) ^14^. All F_0_ were identified as *An. funestus* using *Anopheles funestus* cocktail PCR ^57^ and the resistance profiles of the populations were as reported in the cited articles. Two *An. funestus* laboratory colonies were also used in this study: the FANG colony, which is a fully insecticide-susceptible colony originating from Angola and maintained in laboratory conditions, and the FUMOZ colony, which is a multi-insecticide-resistant colony originating from southern Mozambique, all maintained under laboratory conditions since early 2000 ^51^. The above mosquitoes have been subjected to insecticide bioassays, with their resistance profiles published in the above studies. The female mosquitoes alive 24 h post-exposure were stored in −80°C and were used for this study.

### Africa-wide polymorphism analysis of *An. funestus CYP6P4a* and *CYP6P4b*

Amplification and cloning of full-length cDNA of *An. funestus CYP6P4a* and *CYP6P4b*: To investigate the patterns of genetic variability and potential signatures of selection of *CYP6P4a* and *CYP6P4b* in *An. funestus* across Africa, the full-lengths of *CYP6P4a* and *CYP6P4b* were amplified from pyrethroid-resistant female mosquitoes. Total RNA from pools of 10 permethrin-resistant mosquitoes from the aforementioned origins was extracted using the PicoPure RNA isolation kit (Arcturus, Applied Biosystems, Waltham, Massachusetts, USA). The purified RNA was used for cDNA synthesis using SuperScript III (Invitrogen) with oligo-dT20 and RNase H (New England Biolabs Massachusetts, USA). The full length of the alleles of both genes were then amplified from the cDNA using Phusion Taq polymerase under the following conditions: 1 cycle at 98°C for 1 min; 35 cycles of 98°C for 10 s; 60°C for 30 s; and 72°C for 1 min and 20 s; and 1 cycle at 72°C for 10 min. The primers used are listed in Table S8. Amplicons were gel purified using the QIAquick® Gel Extraction Kit (QIAGEN, Hilden, Germany), and using the CloneJET PCR Cloning Kit, purified products were ligated to the Thermo Ficher Scientific® pJET1.2/blunt cloning vector (Thermo Ficher Scientific Waltham, Massachusetts, USA) and cloned in *E. coli DH5α* competent cells. Minipreparations of plasmids was done using the QIAprep® Spin Miniprep Kit (QIAGEN, Hilden, Germany) and sequenced on both strands using pJET1.2-specific primers (Microsynth AG, Switzerland). Nucleotide polymorphisms were identified through manual examination of sequences and multiple sequence alignments using BioEdit 7.0.5 ^54^. Population genetics parameters of polymorphism like nucleotide diversity π, haplotype diversity, and Tajima’s D and Fu and Li Tajima D* selection estimates of both genes were determined using DnaSP 5.1 software ^53^. Different haplotypes of the genes were compared by constructing a maximum likelihood phylogenetic tree using MEGAX ^52^. The best-fit substitution model based on the Bayesian information criterion that best described the haplotype dataset was the Jones-Taylor-Thornton Gamma distribution model. This was then used to generate the maximum likelihood tree with 1000 bootstrap replications to assess the robustness of the tree. Additionally, haplotype networks were constructed using the TCS program ^58^ to assess the connection between the various haplotypes and pyrethroid resistance.

### Assessment of the temporal variation of the *CYP6P4a* and *CYP6P4b* resistance alleles in Africa

In order to evaluate the temporal dynamics of *CYP6P4a* and *CYP6P4b* between 2014 and 2021, Polymorphism pattern of the *CYP6P4a and CYP6P4b* genes was analysed across Africa using a previously generated SureSelect dataset, which included samples collected in 2014 from Uganda, Malawi, and Cameroon. *CYP6P4a and CYP6P4b* polymorphisms were extracted from the SNP Multisample report file generated through Strand NGS 3.4 for each population. Bioedit ^54^ was utilised to introduce different polymorphisms into the VectorBase reference sequence while ambiguity codes were employed to indicate heterozygote positions in the sequences. Samples analysed include Malawi, Uganda, Cameroon, FANG, and FUMOZ. For the Ghana population, the genes from permethrin resistant mosquitoes from a 2014 collection were cloned and sequenced. Haplotype reconstruction and polymorphism analyses were conducted using DnaSP 5.1 ^53^ and MEGAX ^52^ was used to construct the maximum likelihood phylogenetic trees for both *CYP6P4a* and *CYP6P4b* genes.

### Assessment of copy number variation of C*YP6P4a* in Ghana

Genome-wide analyses of the *An. funestus* population of Ghana revealed the presence of a 6.2kb insertion between *CYP6P4a* and *CYP6P5* ^31^. Quantitative PCR (qPCR) was used to assess whether a potential copy-number variation or gene amplification of *CYP6P4a* could be associated with its significant up-regulation in resistant mosquitoes. Genomic DNA (gDNA) from Ghana mosquitoes and FANG were extracted ^59^. The gDNA concentration and integrity were assessed using an Implen nanophotometer N50 (Implen, Munich, Germany). The concentrations were adjusted to 14ng/µL, and 1µL of gDNA was used in a qPCR reaction for the quantification of *CYP6P4a* (Primers in Table S8). The fold change of the gene in Ghana compared to FANG was calculated using the 2^-ΔΔCT^ equation ^60^ after normalization with housekeeping genes Actin (AFUN006819-RA) and RSP7 (ribosomal protein S7; AFUN007153-RA).

### *In silico* characterisation of CYP6P4a and CYP6P4b

#### Homology modelling of protein structures

The 3-D atomic resolution structures of the major resistant alleles identified in Ghana (CYP6P4a*-*GHA) and one from FANG (CYP6P4a-FANG) were modelled using AlphaFold ^20^, with the default settings implemented in Python 3 running on T4 GPU system. Of three retained models predicted for each allele, models with the highest average estimated reliability score (pLDDT: predicted local distance difference test) were selected for downstream simulations. In addition, model quality was assessed against the crystal structure of human microsomal P450, *CYP3A4*(PDB 1TQN) as a template ^61^, with 33.82% and 33.40% identity to CYP6P4a-GHA and CYP6P4a-FANG query sequences respectively. The ligand structures of deltamethrin (PubChem CID: 40585) and permethrin (PubChem CID: 40326) were retrieved from the PubChem database (https://pubchem.ncbi.nlm.nih.gov/).

#### Structure preparation and suitability for docking

Ligand and protein models were prepared for docking using structure preparation modules implemented in the Molecular Operating Environment (MOE 2021) as described previously ^62^. In brief, solvent and ligand atoms were removed from the CYP3A4 template, followed by modelling of missing residues and broken loops (S281-S286). Subsequently, all structures were protonated using default parameters, partial charges calculated for each atom, and energy minimised with gas phase parameters using the OPLS-AA ^63^ all atom forcefield with a gradient descent threshold of 0.001 kcal/mol. Structures of the heme ligand and corresponding binding site residues were mapped onto predicted models. Post-structural re-examinations, prepared ligand and protein files were converted to multiple formats (.pdb, .smi, .sdf) for docking.

#### Molecular docking of *CYP6P4a* and *CYP6P4b* with pyrethroids

The Dock module implemented in MOE was used for docking computations. First, the docking protocol was optimised and validated by varying pairs of scoring function parameters. This was achieved by re-docking the native heme molecule with its apo CYP3A4 structure and to heme-mapped binding pockets of predicted models, aiming at reproducing the original heme binding pose. The scoring function pair with optimal performance was retained and include Affinity dG for preinitial-scoring, and the Generalized-Born Volume Integral/Weighted Surface Area (GBVI/WSA) for rescoring ^64^. GBVI/WSA is an AMBER99 forcefield ^65^ parameterised function trained on 99 experimentally determined protein-ligand complexes and known to estimate binding free energies of complexes with great accuracy. The retained scoring functions consistently re-produced heme binding poses with negligible deviations (RMSD < 0.5Å) between predicted and experimental binding modes. Secondly, optimised parameters were applied to dock insecticides separately with the respective targets. Substrate recognition site (SRS) residues above the heme catalytic centre were used to define the binding pockets of the models and include: SRS1(H121, F123, A124), SRS2(F213), SRS4(F310, V311, L314), SRS5 (I381), SRS6(S494, F495). In addition, the Ghana mutations L406F; V407I; L409I were also included in binding site due to proximity. A 250ns thermodynamic Integration simulation was subsequently performed to validate molecular docking results. Structural analyses and visualisation were performed with the PyMOL and MOE.

### Heterologous expression of alleles in *E. coli* and insecticide metabolism assay

#### Cloning of *CYP6P4a* and *CYP6P4b* candidate alleles

Candidate CYP6P4a and CYP6P4b alleles were cloned and expressed in *E. coli* cells and used for insecticide metabolism assays. The expression constructs were designed as described in previous work ^66^. Briefly, expression plasmids were constructed (primers in Table S8) by fusing cDNA fragment from a bacterial ompA+2 leader sequence with its downstream ala-pro linker to the NH_2_-terminus of the P450 cDNA, in frame with the P450 initiation codon, as described in previous studies ^25,29,66^. These constructs were digested with the *NgoM*IV and *Xba*I restriction enzymes and ligated to pCW-ori+ expression vector, linearised with the same restriction enzymes, creating the constructs pB13::OMPA+2-CYP6P4a for CYP6P4a sequences (CYP6P4a-GHA, CYP6P4a-FANG, and CYP6P4a-MWI, for Ghana, FANG, and Malawi alleles, respectively). Similarly, the constructs pB13::OMPA+2-CYP6P4b were created for *CYP6P4b* sequences (CYP6P4b-GHA, CYP6P4b-FANG, CYP6P4b-MOZ, and CYP6P4b-UGA, for Ghana, FANG, Mozambique, and Uganda alleles, respectively).

#### Heterologous expression of recombinant *CYP6P4a* and *CYP6P4b*

The *E. coli JM109* cells were co-transformed with the above P450 constructs and a plasmid containing the *An. gambiae* cytochrome P450 reductase (pACYC-AgCPR). Cultured cells in LB medium were allowed to grow until reaching an optical density at 600nm of 0.7-0.8 before addition of the heme precursor δ-aminolevulinic acid (ALA), to a final concentration of 0.5 mM and isopropyl-1-thio-β-D-galactopyranoside (IPTG) to a final concentration of 1 mM were added. Membranes were isolated as done previously and P450 contents determined using spectral analysis ^67^, while CPR activities were conducted following established protocols as previously described ^25,38^. Briefly, about 22 h post-induction, cells were harvested and spheroplasts were prepared and sonicated. The membrane fractions containing P450s were then isolated by ultracentrifugation at 50,000g and resuspended in TSE buffer (50 mM Tris, pH 7.6, 250 mM sucrose, 10% glycerol), and stored in −80°C following measurement of P450 contents.

#### Insecticide metabolism assays and HPLC analysis

The Insecticide metabolism assay was carried out following the method previously described ^25^. Briefly, the following were added to 1.5 ml tubes chilled on ice: 0.1µM of purified P450, 0.025 M potassium phosphate at pH 7.4, 0.25 mM MgCl_2_, 1 mM glucose-6-phosphate, 1 unit/mL glucose-6-phosphate dehydrogenase (G6PDH), 0.8 µM cytochrome b_5_, and 0.2 mM of test pyrethroid insecticide. The tubes were preincubated at 30°C and 1200 rpm for 5 min to activate membranes. Then, 0.1 mM NADP was added to the tubes in a final volume of 200 ml. Reactions were started and carried out at 30°C and 1200 rpm for 90 mins. All reactions were carried out in triplicate, with test reactions containing NADPH (NADPH+) and negative controls lacking NADPH (NADPH-). After 90 mins incubation, 200 μl of ice-cold acetonitrile was added to the tubes to quench reactions and the tubes incubated for 5 more min. The tubes were centrifuged at 16000g for 20 min at 4 °C, and 150 μl of supernatant were transferred to HPLC vials for the quantification of pyrethroid remaining in the samples using reverse-phase HPLC. Substrate peaks were separated with a 250 mm C18 column (Acclaim 120, Dionex) on an Agilent 1260 Infinity (Agilent, Waldbronn, Germany). Enzyme activity was quantified as percentage depletion of insecticide (difference in the amount of insecticide left) between the test (NADPH+) and control (NADPH-). Student t-test was used for the estimation of significance.

### *In vivo* transgenic expression of *CYP6P4a* and *CYP6P4b* in *Drosophila* melanogaster flies

To investigate if overexpression of *CYP6P4a* and *CYP6P4b* alone can confer pyrethroid resistance to a model organism and establish if the intensity of the resistance is impacted by allelic variation, *CYP6P4a* and *CYP6P4b* were expressed in Drosophila flies using the GAL4/UAS system ^29,68^. Transgenic flies were then used for pyrethroid susceptibility tests.

#### Cloning and construction of transgenic Drosophila lines

PCR amplification of the candidate alleles was done using minipreps that were used for *in vitro* expression above. PCR was carried out using Phusion High-Fidelity DNA Polymerase with primers having the restriction sites for *Eag*I and *Xba*I restriction enzymes (Table S8). PCR and cloning protocols were described in previous studies ^25,29^. The amplicons were purified and cloned in *E. coli DH5α* competent cells using pJET1.2 vector (Thermo Ficher Scientific Waltham, Massachusetts, USA). Clones were digested with the above restriction enzymes, and the digests were ligated to pUASattB vector linearised with the above restriction enzymes. The constructs: *CYP6P4a* (UAS-CYP6P4a-GHA, and UAS-CYP6P4a-FANG) and *CYP6P4b* (UAS-CYP6P4b-GHA, UAS-CYP6P4b-FANG, and UAS-CYP6P4b-MOZ) were cloned into *E. coli DH5α* competent cells, miniprepped, and injected into the germline of *D. melanogaster* line carrying the attP40 docking site, 25C6 on chromosome 2 [y w M (eGFP, vas-int, dmRFP) ZH-2A; P{CaryP} attP40 using the PhiC31 integrase system. The process of microinjection and balancing of UAS stock to eliminate the integrase was conducted by the Fly Facility (Cambridge, UK), generating the respective transgenic lines. Ubiquitous expression of candidate alleles was obtained by crossing Actin5C-GAL4 (GAL4-Actin driver strain Act5C-GAL4, BL25374 [y[1] w[*]; P{Act5C-GAL4-w}E1/CyO, 1;2], Bloomington, IN, USA) virgin female flies with transgenic homozygote UASCYP6P4a and UAS-CYP6P4b males to produce F_1_ experimental progeny (Act5C-CYP6P4a and Act5C-CYP6P4b). For the control group, male flies with the same genetic background as the UAS transgenic lines but devoid of the pUASattB-CYP6P4a or pUASattB-CYP6P4b constructs were crossed with virgin females from the Actin5C-GAL4 driver line to generate Actin5C-GAL4-null flies.

#### Insecticide susceptibility contact assays

Five replicates of F_1_ female flies (2-5 days old) from experimental and control groups were exposed to pyrethroids (4% permethrin-, 0.2% deltamethrin-, and 0.0007% alphacypermethrin-impregnated filter papers prepared in acetone and Dow Corning 556 Silicone Fluid (BHD/Merck, Germany), with mortality plus knockdown recorded after 1, 2, 3, 6, 12, and 24 h. A comparison of the mortality rates between the experimental groups and the control groups was used to assess if candidate genes’ overexpression alone is enough to confer resistance, while a comparison with flies harbouring the FANG allele shed light on the impact of allelic variation on resistance.

To confirm the overexpression of candidate genes, qRT-PCR was carried out using RNA extracted from three replicates of pools of five experimental and control female flies. One microgram of total RNA from each of the three biological replicates of the experimental and control groups was used for cDNA synthesis using SuperScript III (Invitrogen) with oligo-dT20 and RNase H, according to the manufacturer’s instructions. A serial dilution of cDNA was used to establish standard curves for each gene to assess PCR efficiency and quantitative differences between samples. The quantitative PCR (qPCR) amplification was carried out in a MX 3005 real-time PCR system (Agilent) using Brilliant III Ultra-Fast SYBR Green QPCR Master Mix (Agilent). A total of 10 ng of cDNA from each sample was used as template in a three-step program involving a denaturation at 95 °C for 3 min followed by 40 cycles of 10 s at 95 °C and 10 s at 60 °C and a last step of 1 min at 95 °C, 30 s at 55 °C, and 30 s at 95 °C. The relative expression and fold-change of each target gene in R and C relative to S was calculated according to the 2^−ΔΔCT^ method incorporating PCR efficiency (14) after normalization with the housekeeping RSP7 ribosomal protein S7 (AGAP010592) and the actin 5C (AGAP000651) genes A quantitative reverse transcription PCR reaction was carried out with normalization with the Actin2 and RSP7 housekeeping genes using primers. All primers are provided in Table S8.

#### Design of DNA diagnostic marker assays for *CYP6P4a* and *CYP6P4b* mutations

A single, most predominant mutation identified in the Ghana-specific variants of *CYP6P4b* (D284E) and *CYP6P4a* (M220I) were targeted to create DNA-based diagnostic assays (CYP6P4b-D284E and CYP6P4a-M220I). For *CYP6P4b*, the design exploited the Amplification Refractory Mutation System (ARMS) PCR principle http://primer1.soton.ac.uk/primer1.html, that allowed the specific amplification of the wild-type allele (FANG) and the mutant allele (Ghana). Briefly, a set of four primers was designed with two outer primers: 6P4b_ARMS_OF: 5′-CTA TCT GCT CAT CTG TTT GCA CTG GA-3′ and 6P4b_ARMS_OR: 5′-GTA TGT CCG TTC TGC ACC C-3′, and two allele specific inner primers: 6P4b_ARMS_CF: 5′-CTG CAG ATT AAG AAC AAA GGT TAT TTG AAC-3′ and 6P4b_ARMS_AR: 5′-TTA TCA TTG GCT CCA ATG TCA CGT TTT T-3′ (Integrated DNA Technology, Belgium). A PCR of 35 cycles was carried out using the four primers using the conditions described in Table S4. Distinct genotype amplicons were visualised using 1.5% agarose gel electrophoresis. The homozygous wild-type allele had the S genotype band at 809 bp and the common band at 1284 bp. The homozygous mutant had the R genotype band at 527 bp and the common band at 1284 bp. The heterozygous genotype had three band comprising 527 bp, 809 bp and 1284 bp. For *CYP6P4a,* a Locked Nucleic Acid (LNA) probe-based PCR (Integrated DNA technologies, UK) was designed to discriminate between the wild-type and mutant alleles of *CYP6P4a.* The design comprises two primers: 6P4a_F 5′-ATAC GGC AAC AAG GTG TTC-3′ and 6p4a_R 5′-CCT TCG TCA GTC AGC TTA AC-3′ and two probes: the wildtype specific probe LNA6p4a-Met: Hex: TGT+TCTTA+T+G+GT+AA+A+GT and the mutant-specific probe LNA6p4a-Ile:Fam: ACTGT+C+CTTAT+T+TT+C+AA+AT (Integrated DNA Technology, Belgium). The PCR involves 10 minutes denaturation at 95°C; (Segment 1) and 40 cycles of denaturation for 10 seconds at 95°C, annealing for 45 seconds at 60°C; (Segment 2) (see Table S5 for PCR details). Distinct genotypes are identified with different fluorophores using the MxPro software https://www.agilent.com/en/product/real-time-pcr-(qpcr)/real-time-pcr-(qpcr)-instruments/mx3000-mx3005p-real-time-pcr-system-software. Detailed information on the PCR reactions can be found in Table S5.

#### Association between the *CYP6P4a* and *CYP6P4b* markers and pyrethroid resistance

The high level of pyrethroid resistance, coupled with the high frequency of the resistance alleles in the Ghanaian field population made the establishment of the link between the genotype and the resistance phenotype difficult. To address this challenge, a mosquito genetic cross between female FANG bearing the homozygote susceptible (SS) CYP6P4b D284/D284 and CYP6P4a M220/M220 genotypes and male field mosquitoes from Ghana bearing the homozygote resistant (RR) CYP6P4b E284/E284 and CYP6P4a I220/I220 genotypes was established as done for the development of *CYP6P9a*/b markers ^14,15^. The crosses were maintained through to the fourth generation to allow the three genotypes (RR, RS, and SS) to segregate. WHO tube bioassays were carried out on three- to five-day-old hybrid female mosquitoes using pyrethroid insecticide papers following WHO protocol ^5^ and the correlation between resistance phenotype and genotypes was established using odds ratio.

#### Impact of *CYP6P4a* and *CYP6P4b* resistance alleles on the bio-efficacy of bed nets

The effectiveness of current pyrethroid-treated bed nets in relation to the impact of the *CYP6P4a* and *CYP6P4b* resistant alleles was assessed using cone bioassays ^69^. Bed nets evaluated were Olyset Net (1000 mg/m^2^ permethrin), PermaNet 2.0 (55 mg/m^2^ deltamethrin), and DuraNet (261 mg/m^2^ alpha-cypermethrin). Five replicates of ten females (2- to 5-d old) from F_3_ and F_4_ generations were placed in plastic cones enclosed with the aforementioned bed nets for 3 mins. Post-exposure to the bed nets, mosquitoes were transferred to holding paper cups and fed with cotton soaked in a 10% sugar solution. Final mortality rates were assessed 24 h later, and DNA was extracted from 30–40 alive and dead mosquitoes using the LIVAK protocol, and the genotypes were determined. Mortality and the correlation between resistance phenotype and *CYP6P4a* and *CYP6P4b* genotypes were used to evaluate the efficacy of the bed nets.

### Africa-wide distribution of *CYP6P4a* and *CYP6P4b* resistance alleles

The Africa-wide geographical distribution of the *CYP6P4a* and *CYP6P4b* resistance alleles was assessed by genotyping genomic DNA samples of female *An. funestus* collected in different regions of Africa in different years. Central: Cameroon-Mibellon (2021); Southern: Malawi-Chikwawa (2021); Eastern: Uganda-Mayuge (2022), Tanzania-Maheza (2017), 38° 54’ 14.9904’’ E; https://www.countrycoordinate.com/ and Western Africa: Benin (2022), Seirra-Leone-Largo (2021), 12° 8’ 56.32”W https://www.longitude-latitude-maps.com/city/.

Thirty to forty parental mosquitoes were genotyped per locality using the above diagnostic tools.

### Statistical analysis

Statistical analysis was done using GraphPad Prism 7. Statistical significance was set at 0.05 and 95% confidence interval (CI). Comparison of the insecticide depletion of enzymes and transgenic insecticide contact bioassay was done using student t-test. The correlation between the CYP6P4a_R and the CYP6P4b_R markers and their impact on pyrethroid resistance phenotype and on the bio-efficacy of bed nets was established using odds ratio (OR) and Fisher’s exact test.

## Acknowledgements

We thank the staff of the Centre for Research in Infectious Diseases (CRID) for their support and assistance throughout this study. Special appreciation goes to Murielle Wondji and Helen Irving for handling the sequencing processes and laboratory needs. A special thanks to Dr Mark Paine (LSTM) for providing the technical platform for the *in vitro* enzyme characterisation, and we will also like to extend gratitude to Prof. Mark R. Sanderson for reviewing the *in silico* characterisation of this study. Finally, we want to thank all the inhabitants of the different African sample collection sites, especially those of Atatam in Ghana for their generous support during the study.

## Author contributions

Conceptualisation: CSW

Methodology: NMTT-N LMJM, DNN, AM, TK, SSI, CSW

Investigation: NMTT-N, LMJM, AM, DNN, MFMK, CJT, MT, SSI, CSW

Supervision: CSW, SGM

Writing—original draft: NMTT-N, DNN, and CSW

Writing—review & editing: NMTT-N, LMJM, AM, DNN, MFMK, MT, SGM, SSI, CSW

### Consent for publication

All authors declare consent for publication.

