## Supplementary material for "Two highly selected mutations in the tandemly duplicated *CYP6P4a* and *CYP6P4b* drive pyrethroid resistance in *Anopheles funestus*": Supplemetary file

### Supplementary Figures

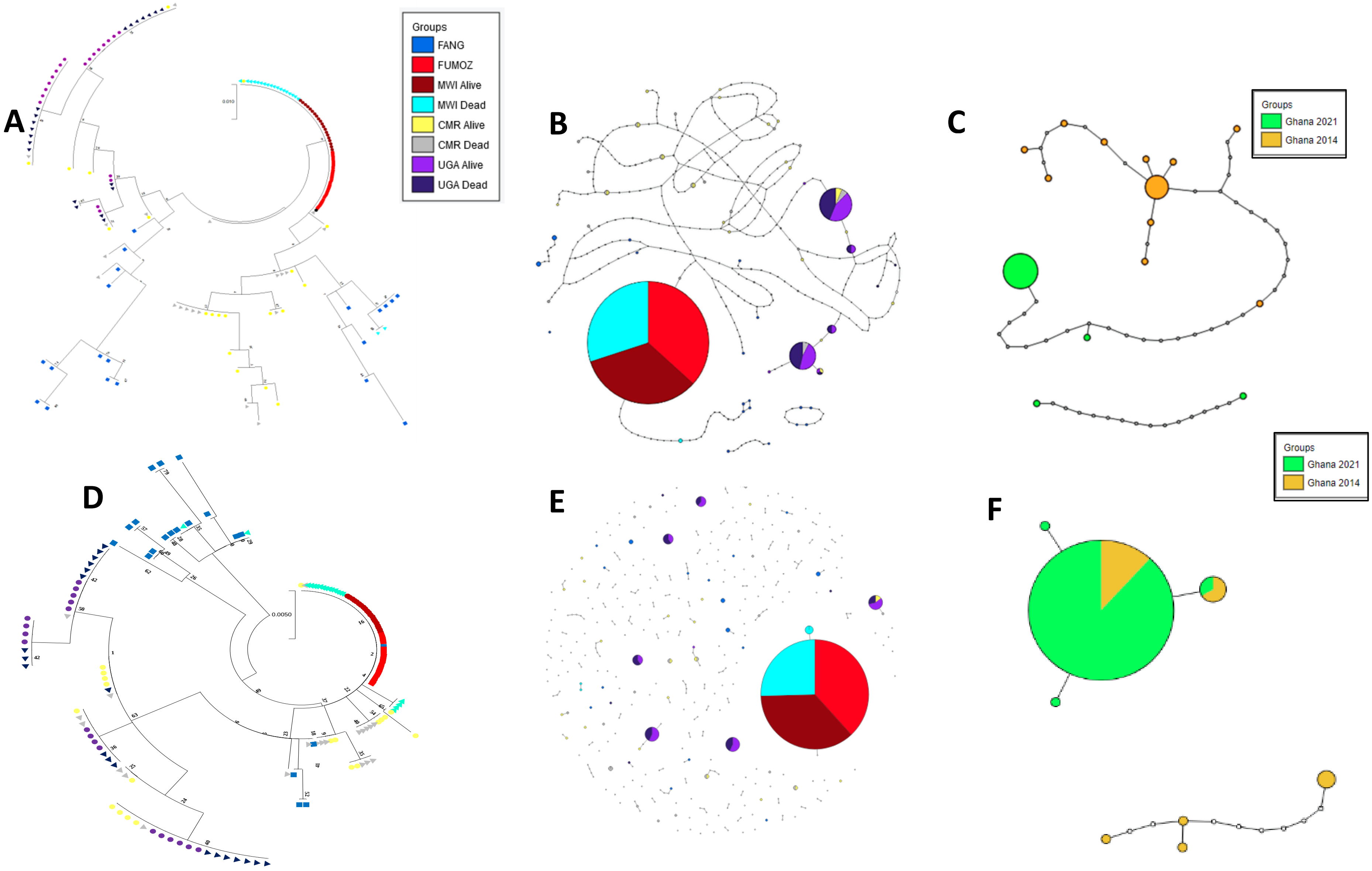

**Fig. S1. Polymorphism patterns of *CYP6P4a* and *CYP6P4b* from 2014 sureselect and sanger sequencing data. A.** Phylogenetic tree for *CYP6P4a* from 2014 sureselect data across Cameroon, Uganda, Malawi*;* **B**. TCS haplotype network for *CYP6P4a* from 2014 sureselect data across Cameroon, Uganda, Malawi. A fixed haplotype is observed in Southern population (Malawi and FUMOZ) and the formation of major haplotypes in Uganda, but high diversity in Cameroon; **C.** TCS haplotype network for *CYP6P4a* from Ghana 2014 and 2021 samples showing high diversity in 2014 but selection of major resistance haplotypes in 2021; **D**. Phylogenetic tree for *CYP6P4b* from 2014 sureselect data across Cameroon, Uganda, Malawi*;* **E.** Haplotype network for *CYP6P4b* from 2014 sureselect data across Cameroon, Uganda, Malawi. Similarly, A fixed haplotype is observed in Southern population and the formation of major haplotypes in Uganda, but very high diversity in Cameroon; **F**. Haplotype network for *CYP6P4b* from Ghana 2014 and 2021 samples showing high diversity in 2014 but with the presence of resistance alleles that have undergone selection to near fixation in 2021.

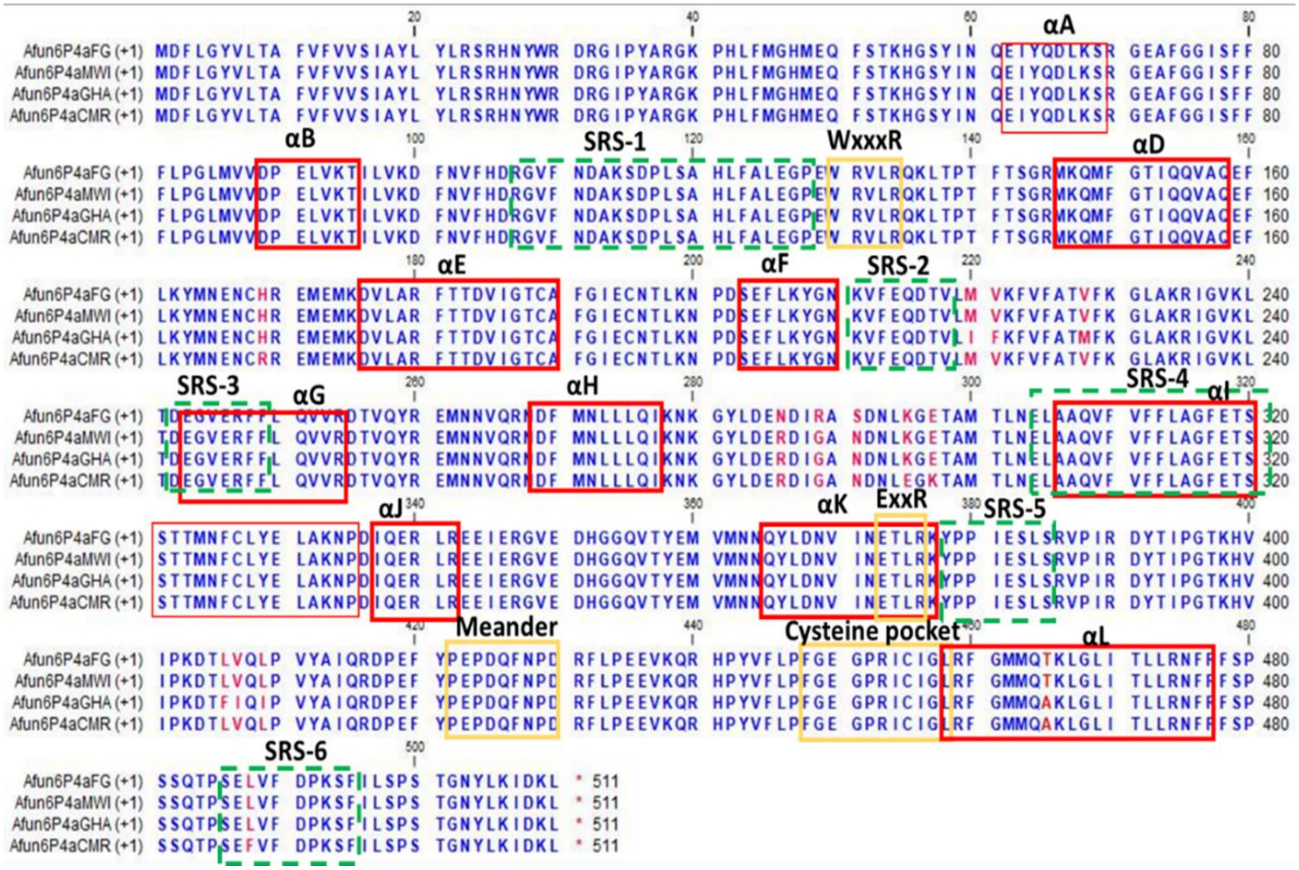


**Fig. S2A. Amino-acid alignment of CYP6P4b variants highlighting substitutions.** Key conserved P450 motifs and substrate recognition sites are annotated. The solid red lines represent helices A-L, while dashed blue lines correspond with the substrate recognition sites 1-6. Solid orange lines identify the structurally conserved motifs of the CYP450s. Variable residues are in hot pink. Residues 310-315 corresponds to the oxygen binding pocket.

A

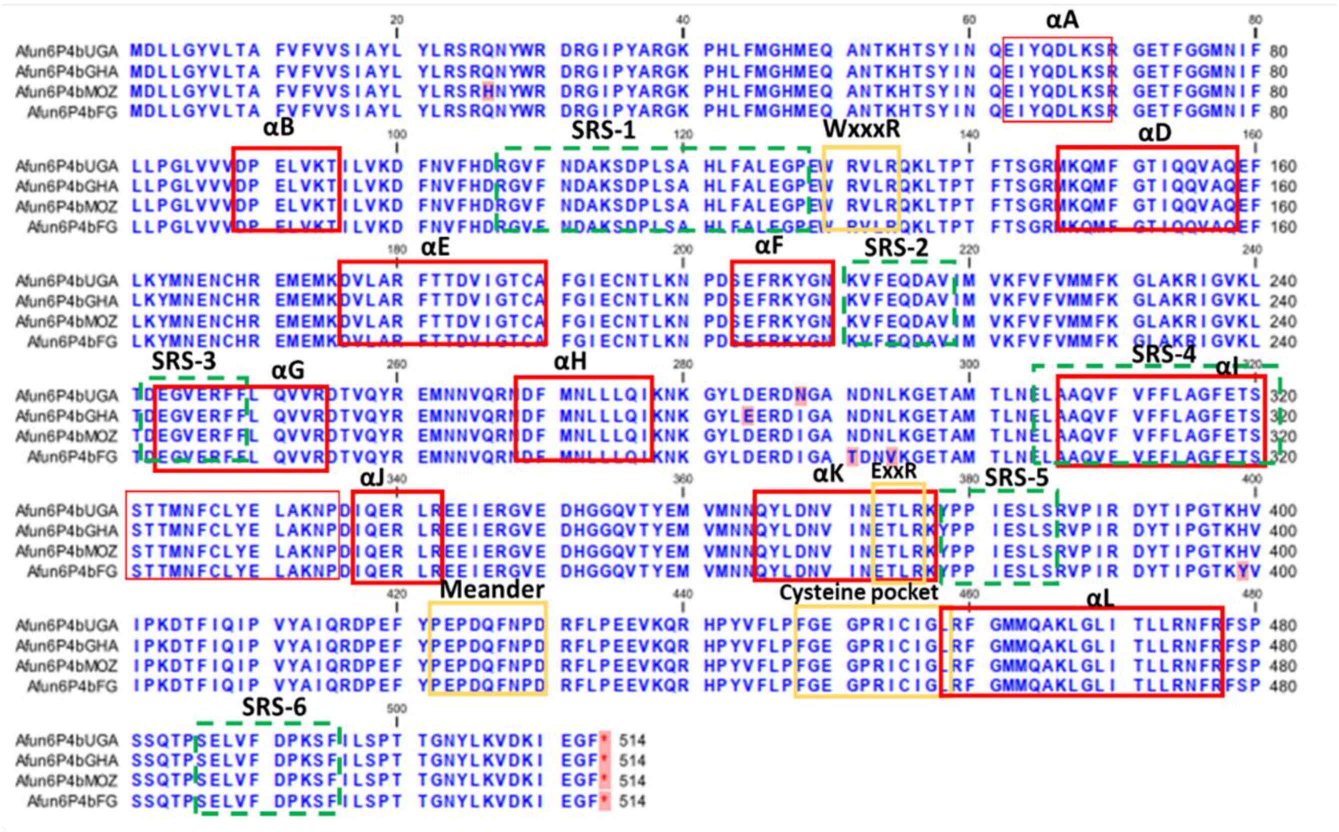

B

| Hap | Nb | 26 | 29 | 30 | 61 | 86 | 264 | 284 | 288 | 291 | 294 | 399 |
|---|---|---|---|---|---|---|---|---|---|---|---|---|
| FUMOZ/MOZ | 16 | H | W | R | Q | V | N | D | I | N | L | H |
| GHA 2021 | 16 | Q | . | . | . | . | . | E | . | . | . | . |
| GHA 2021 | 1 | Q | . | . | . | . | D | E | . | . | . | . |
| GHA 2021 | 1 | Q | . | . | R | . | . | E | . | . | . | . |
| GHA 2014 | 4 | Q | . | . | . | F | . | . | . | . | . | . |
| GHA 2014 | 4 | Q | . | . | . | . | . | E | . | . | . | . |
| GHA 2014 | 1 | Q | . | . | . | F | . | . | . | . | . | . |
| CMR | 5 | Q | . | . | . | . | . | . | N | . | . | . |
| CMR | 1 | Q | . | C | . | . | . | . | N | . | . | . |
| CMR | 1 | Q | R | . | . | . | . | . | N | . | . | . |
| UGA | 10 | Q | . | . | . | . | . | . | N | . | . | . |
| FANG | 3 | Q | . | . | . | . | . | . | . | T | V | Y |
| FANG | 7 | Q | . | . | . | . | . | . | . | . | . | . |
| FANG | 1 | Q | . | . | . | . | . | . | . | T | V | . |

**Fig. S2B. A. Amino-acid alignment of *CYP6P4b* variants highlighting substitutions.** Key conserved P450 motifs and substrate recognition sites are annotated. The solid red lines represent helices A-L, while dashed blue lines correspond with the substrate recognition sites 1-6. Solid orange lines identify the structurally conserved motifs of the CYP450s. Variable residues are in hot pink. Residues 310-315 corresponds to the oxygen binding pocket. **B.** Schematic representation of haplotypes of *CYP6P4b* with D284E mutation occurring in Ghana highlighted in red.

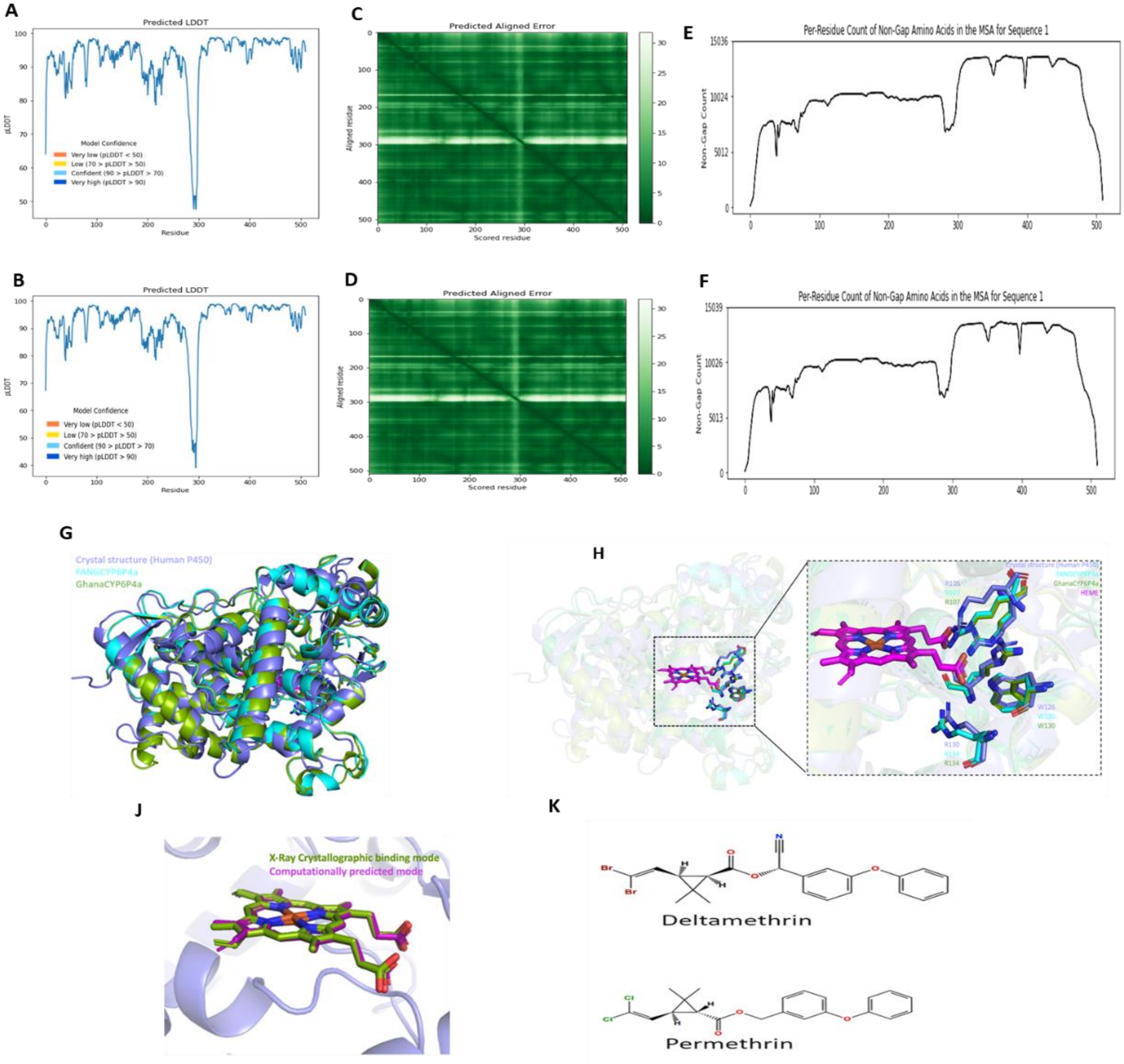


**Fig. S3. AlphaFold's predicted per residue accuracy (pLDDT) of modelled structures that estimates the confidence level of predictions.** A. CYP6P4a-GHA and B. CYP6P4a-FANG. The Predicted aligned error reflects the relative structural prediction confidence of different domains in the structures C. CYP6P4a-GHA D. CYP6P4a-FANG. Multiple sequence alignments (MSA) with AlphaFold's big fantastic database (BFD) for E. CYP6P4a-GHA and F. CYP6P4a-FANG. G. Structural alignment of CYP6P4a-GHA (colored green) and CYP6P4a-FANG (colored cyan) alleles, onto the crystal structure of human microsomal cytochrome P450 (colored slate blue; PDB 1TQN). H. Highlight of heme pocket showing optimal alignment (RMSD<0.37Å) between heme (colored purple) interacting side chain residues of the three models. J. Validation of docking protocol: heme molecule (colored green) from the X-ray experimental structure was re-docked into its binding pocket. The software was most accurately able to predict a heme binding mode (colored purple pink) very closely (RMSD=0.2Å) to the crystal one. K. 2D-structures of Deltamethrin and Permethrin

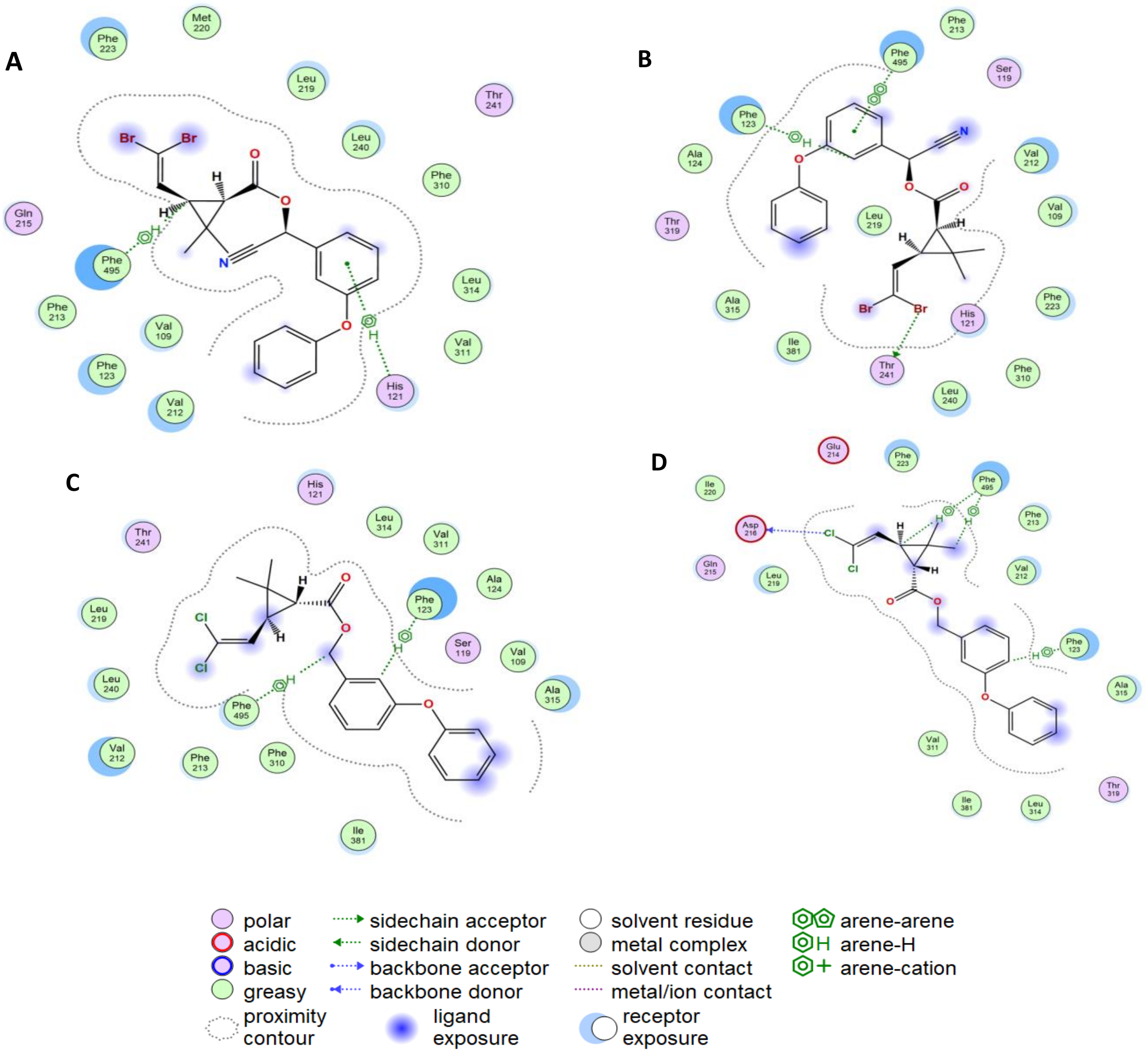


**Fig. S4.** 2-D interaction map of representative pyrethroid poses with 4'-phenoxy spot approaching above the heme iron at a distance ranging between 1.5 - 6.5Å. Deltamethrin bound to **A.** FANGCYP6P4a and **B.** GhanaCYP6P4a. 2-D interaction map of Permethrin bound to **C.** FANGCYP6P4a and **D.** GhanaCYP6P4a. Contour lines, arene-arene, arene-H, arene-cation are all estimations of van der Waals interactions. The closer the contour line to the pyrethroid, the higher the vdW clash.

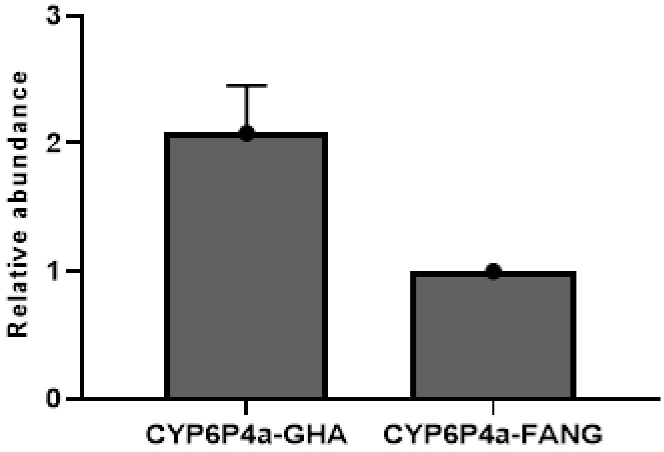


**Fig. S5. Genomic abundance of *CYP6P4a* in Ghana samples relative to abundance in FANG. Data** confirms a duplication of the gene in the Ghanian mosquitoes

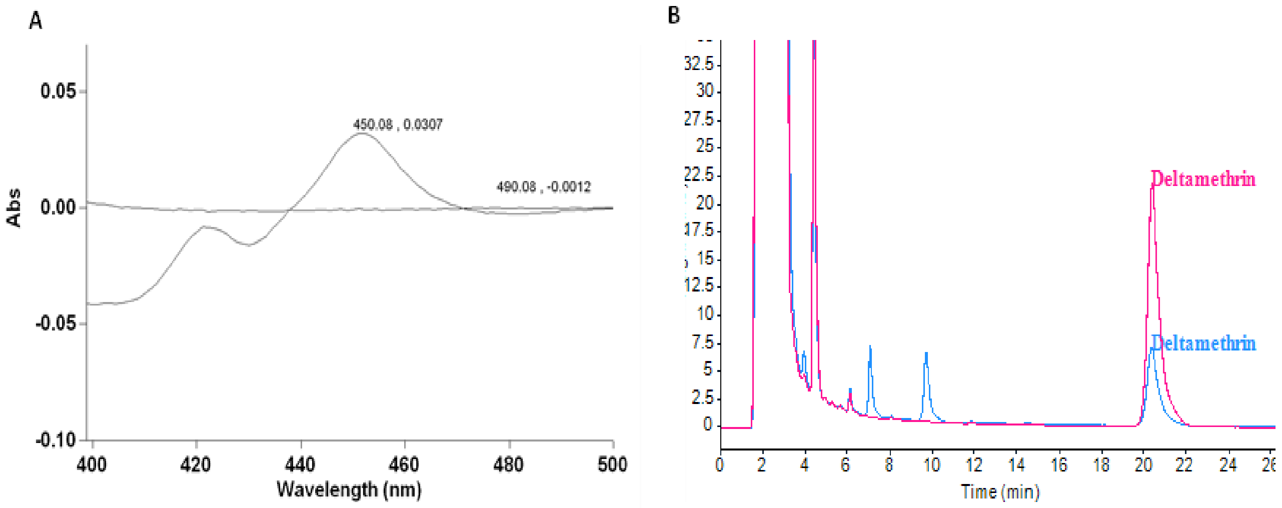


**Fig. S6.** Heterologous expression of candidate alleles and metabolism of insecticides by recombinant CYP6P4a and CYP6P4b enzymes. **A**. CO-difference spectrum generated from E. coli membranes expressing CYP6P4a and CYP6P4b alleles**.** Among the different allelic variants, the FANG variants *6P4a-*$N^{286}R^{289}S^{291}$ (6P4a-FANG) and *6P4b-*$T^{291}V^{294}Y^{399}$ (6P4b-FANG) demonstrated the highest expression levels, producing 15.3 µM and 7.2 µM CYP6P4a and CYP6P4b proteins, respectively. The other resistant variants produced 2-5 µM of recombinant enzymes. **B**. Overlay of HPLC chromatogram of the CYP6P4a and CYP6P4b depletion of deltamethrin with –NADPH (negative control) in hot pink and +NADPH (experimental) in blue.

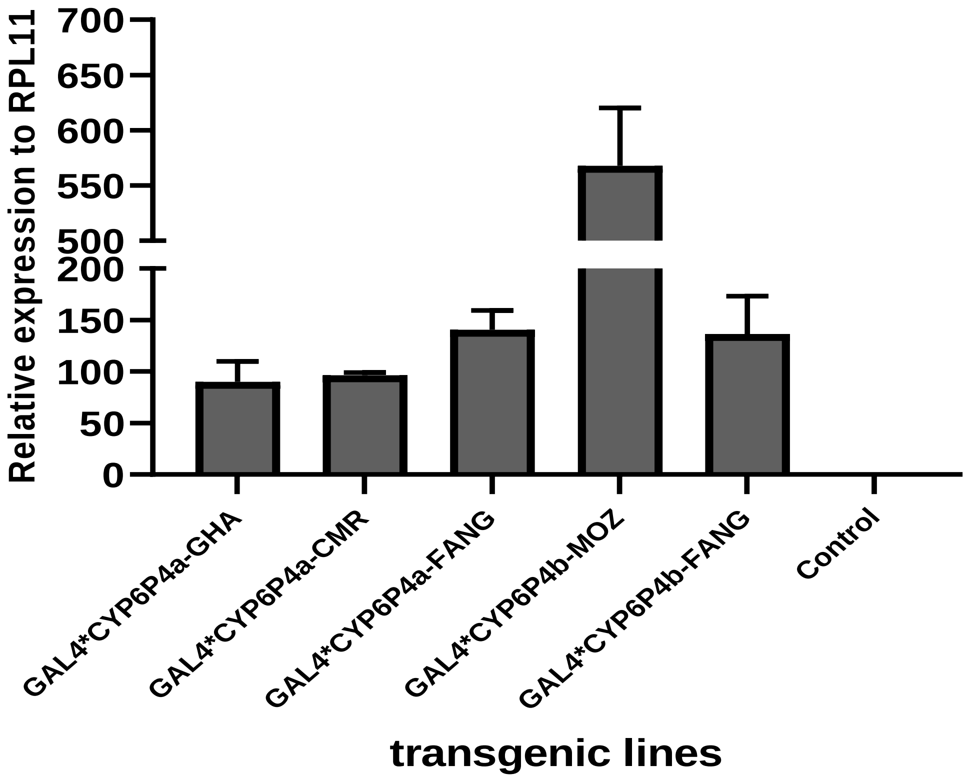


**Fig. S7. Confirmation of transgene expression in *Drosophila melanogaster***

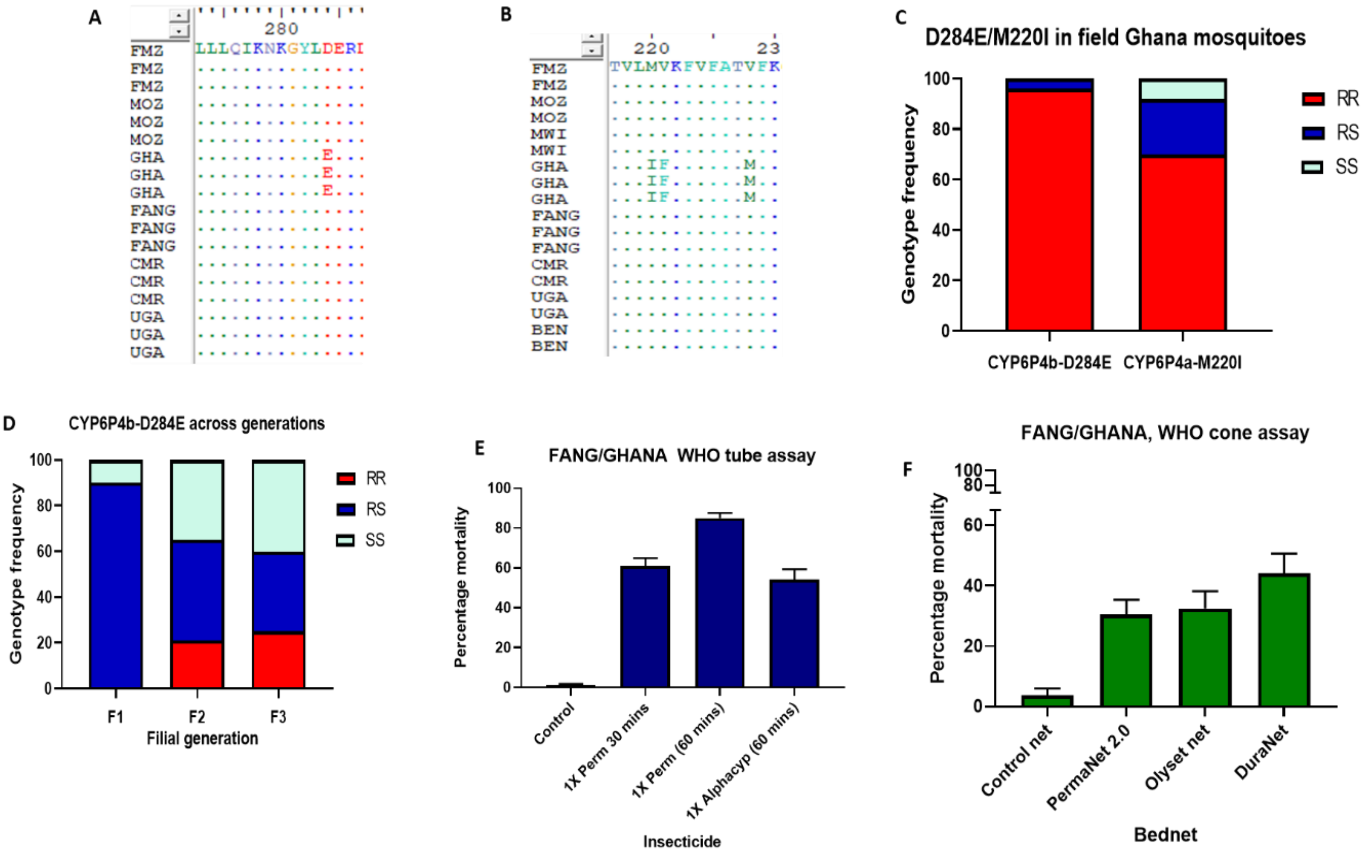


**Fig. S8. Schematic alignment of sequences across Africa. A**. D284E mutation in Ghana and its absence in the other populations, used in the design of the CYP6P4b-D284E molecular diagnostic tool, **B.** M220I mutation in Ghana and its absence in the other populations, used in the design of the CYP6P4a-M220I molecular diagnostic tool. **C.** Frequency of resistance genotypes in Ghanaian population. **D**. Distribution of the CYP6P4b-D284E mutation in the hybrid FANG/GHANA strain showing segregation of genotypes. Susceptibility profile of FANG/GHANA strain to evaluate impact of CYP6P4a-M220I and CYP6P4b-D284E markers on resistance and on bio-efficacy of ITN. **E**. WHO tube assay **F**. WHO cone assay.

#### Supplementary Tables

##### Table S1. Nucleotide diversity parameters of the coding regions of *CYP6P4a* and *CYP6P4b* across Africa.

| Sample | N | S | h | Hd | Syn | Nsyn | π | D | D* |
|---|---|---|---|---|---|---|---|---|---|
| | | | | | | CYP6P4a | | | |
| **Cameroon 2021** | 12 | 0 | 1 | 0 | 0 | 0 | 0 | 0 | 0 |
| **Ghana 2021** | 9 | 38 | 4 | 0.583 | n.a | 11 | 0.009 | -0.013 ns | 1.104 |
| **Ghana 2014** | 14 | 25 | 11 | 0.934 | n.a | 8 | 0.003 | -1.26 ns | -1.411 ns |
| **Fang** | 6 | 20 | 6 | 1 | 15 | 5 | 0.007 | 1.36 ns | 1.211 |
| **Benin 2021** | 6 | 13 | 3 | 0.733 | 10 | 3 | 0.004 | 0.472 ns | 0.6876 ns |
| **Uganda 2021** | 10 | 0 | 1 | 0 | 0 | 0 | 0 | 0 | 0 |
| **Fumoz** | 15 | 0 | 1 | 0 | 0 | 0 | 0 | 0 | 0 |
| **Moz. 2021** | 11 | 0 | 1 | 0 | 0 | 0 | 0 | 0 | 0 |
| **Malawi 2021** | 7 | 0 | 1 | 0 | 0 | 0 | 0 | 0 | 0 |
| **All** | 93 | 84 | 24 | 0.782 | n.a | n.a | 0.012 | 0.57 | 0.89 |
| | | | | | | CYP6P4b | | | |
| **Cameroon 2021** | 7 | 1 | 2 | 0.286 | 1 | 0 | 0.00019 | -1.006 ns | -1.0488 ns |
| **Ghana 2021** | 18 | 2 | 3 | 0.216 | 0 | 2 | 0.00014 | -1.508 ns | -1.989 ns |
| **Ghana 2014** | 9 | 25 | 6 | 0.917 | 23 | 2 | 0.009 | 2.24 * | 1.588 * |
| **Fang** | 11 | 24 | 6 | 0.855 | 21 | 3 | 0.008 | 1.965 ns | 1.398 * |
| **Uganda 2021** | 10 | 0 | 1 | 0 | 32 | 7 | 0.000 | 0 | 0 |
| **Fumoz** | 7 | 0 | 1 | 0 | 0 | 0 | 0 | 0 | 0 |
| **Malawi 2021** | 9 | 0 | 1 | 0 | 0 | 0 | 0 | 0 | 0 |
| **All** | 77 | 69 | 24 | 0.884 | 54 | 16 | 0.011 | 0.761 | -0.240 ns |

N, number of sequences; S, number of polymorphic sites; Syn, Synonymous mutations; Nsyn, Non-synonymous mutations; h, number of haplotypes; Hd, haplotype diversity; π, nucleotide diversity; D and D* Tajima's and Fu and Li's statistics; ns, not significant; n.a not applicable

**Table S2.** Nucleotide diversity parameters of Africa-wide *An. funestus* populations in 2014

| | | | | | | | | | |
|---|---|---|---|---|---|---|---|---|---|
| | | | | **CYP6P4a (1533bp)** | | | | | |
| **Sample** | **N** | **S** | **h** | **Hd** | **Syn** | **Nsyn** | **π** | **D** | **D*** |
| **FUMOZ** | 20 | 0 | 1 | 0 | 0 | 0 | 0 | 0 | 0 |
| **FANG** | 20 | 59 | 18 | 0.989 | 46 | 14 | 0.0179 | 2.6185** | 1.7389** |
| | | | | **Cameroon** | | | | | |
| **Alive** | 20 | 48 | 20 | 1 | 38 | 10 | 0.0138 | 2.2572* | 1.5975 * |
| **Dead** | 20 | 47 | 20 | 1 | 37 | 10 | 0.0134 | 2.2180 * | 1.5927 ** |
| **All** | 40 | 50 | 35 | 1 | 39 | 11 | 0.0137 | 2.8132 ** | 1.7524 ** |
| | | | | **Uganda** | | | | | |
| **Alive** | 20 | 19 | 7 | 0.8 | 16 | 3 | 0.0058 | 2.4732* | 1.2930 ns |
| **Dead** | 20 | 22 | 7 | 0.8 | 18 | 4 | 0.0063 | 2.0935 * | 1.3562 ns |
| **All** | 40 | 23 | 9 | 0.782 | 19 | 4 | 0.0059 | 2.2528 * | 1.7082** |
| | | | | **Malawi** | | | | | |
| **Alive** | 20 | 0 | 1 | 0 | 0 | 0 | 0 | 0 | 0 |
| **Dead** | 20 | 12 | 2 | 0.189 | 8 | 4 | 0.0016 | -1.1767 ns | 1.4611 * |
| **All** | 40 | 12 | 2 | 0.097 | 0 | 5 | 0.0008 | -1.8035 ns | 1.4751 ns |
| | | | | **All countries** | | | | | |
| **All alive** | 60 | 49 | 26 | 0.867 | 39 | 10 | 0.0127 | 2.871 ** | 1.8367 ** |
| **All dead** | 60 | 58 | 27 | 0.884 | 44 | 15 | 0.0132 | 2.1267 * | 2.0728 ** |
| **All Alive and Dead** | 120 | 61 | 43 | 0.817 | 46 | 15 | 0.0129 | 2.3161 * | 2.1068 ** |
| **Total all** | 160 | 78 | 61 | 0.853 | 60 | 19 | 0.0142 | 1.7492 ns | 2.3320 ** |
| | | | | **CYP6P4b (1542 bp)** | | | | | |
| **FANG** | 20 | 48 | 16 | 0.979 | 40 | 9 | 0.0141 | 2.2307 * | 1.6020 ** |
| **FUMOZ** | 20 | 0 | 1 | 0 | 0 | 0 | 0 | 0 | 0 |
| | | | | **Cameroon** | | | | | |
| **Alive** | 20 | 41 | 20 | 1 | 35 | 6 | 0.0118 | 2.3075 * | 1.6997 ** |
| **Dead** | 20 | 42 | 19 | 0.995 | 37 | 5 | 0.0115 | 1.9613 ns | 1.7027 ** |
| **All** | 40 | 46 | 35 | 0.994 | 41 | 6 | 0.0116 | 2.1951 * | 1.8978 ** |
| | | | | **Uganda** | | | | | |
| **Alive** | 20 | 18 | 7 | 0.879 | 15 | 3 | -0.0057 | 2.7905** | 1.2681 ns |
| **Dead** | 20 | 25 | 10 | 0.926 | 20 | 3 | 0.0063 | 1.955 ns | 1.610 ** |
| **All** | 40 | 4 | 11 | 0.888 | 21 | 3 | 0.0060 | 2.0875 * | 1.4241 * |
| | | | | **Malawi** | | | | | |
| **Alive** | 20 | 0 | 1 | 0 | 0 | 0 | 0 | 0 | 0 |
| **Dead** | 20 | 16 | 4 | 0.489 | 11 | 5 | 0.002 | -1.1706 ns | 1.2105 ns |
| **All** | 40 | 16 | 4 | 0.273 | 11 | 5 | 0.001 | -1.8493 ns | 1.1735 ns |
| | | | | **All countries** | | | | | |
| **All alive** | 69 | 41 | 27 | 0.877 | 40 | 8 | 0.0102 | 2.4679 * | 1.9797 ** |
| **All dead** | 60 | 49 | 33 | 0.937 | 45 | 9 | 0.0111 | 2.0115 ns | 2.0321 ** |
| **All Alive and Dead** | 120 | 54 | 49 | 0.907 | 46 | 10 | 0.0106 | 1.8599 ns | 2.2632 ** |
| **Total all** | 160 | 64 | 65 | 0.879 | 50 | 19 | 0.0115 | 1.5198 ns | 2.4332 ** |

N, number of sequences; S, number of polymorphic sites; Syn, Synonymous mutations; Nsyn, Non-synonymous mutations; h, number of haplotypes; Hd, haplotype diversity; π, nucleotide diversity; D and D* Tajima's and Fu and Li's statistics; ns, not significant; π, n.a not applicable

**Table S3.** Characteristics of productive poses binding with pyrethroids 4'-phenoxy spot approaching above the heme iron at a distance ranging between 1.5 - 6.5Å.

| **Molecular Docking** | | |
|---|---|---|
| **Deltamethrin** | **6P4a-FANG** | **6P4a-GHA** |
| 4'-phenoxy binding mode enrichment (%n) | 56% (N=25) | 71.4% (N=21) |
| Mean binding mode distance to Heme iron (Å ± SD)#* | 4.7 ± 0.38 | 3.2 ± 0.42 |
| Mean GBVI/WSA (ΔG Kcal/mol ± SD) | -34.2 ± 1.3 | -35.5 ± 0.68 |
| Mean binding mode strain energy (Kcal/mol ± SD) | 5.8 ± 1.4 | 6.05 ± 0.7 |
| Mean MM affinity (Kcal/mol ± SD)* | -8.6 ± 0.13 | -9.04 ± 0.11 |
| **Permethrin** | **6P4a-FANG** | **6P4a-GHA** |
| 4'-phenoxy binding mode enrichment (%n) | 39.1% (N=23) | 28% (N=25) |
| Mean binding mode distance to Heme iron (Å ± SD)# | 4.3 ± 0.5 | 3.4 ± 0.6 |
| Mean GBVI/WSA (ΔG Kcal/mol ± SD) | -33.5 ± 0.9 | -34.8 ± 1.2 |
| Mean binding mode strain energy (Kcal/mol ± SD) | 4.0 ± 0.67 | 4.5 ± 0.61 |
| Mean MM affinity (Kcal/mol ± SD)* | -8.17 ± 0.11 | -8.85 ± 0.28 |
| **Thermodynamics integration simulation** | | |
| **Deltamethrin** | **6P4a-FANG** | **6P4a-GHA** |
| ΔG (mmgb) | -8.1524 | -8.2958 |
| ΔQ (mol_charge) | 0 | 0 |
| ΔM (Strain) | 0 | 0 |
| **Permethrin** | **6P4a-FANG** | **6P4a-GHA** |
| ΔG (mmgb) | -7.561 | -7.6933 |
| ΔQ(mol_charge) | 0 | 0 |
| ΔM (Strain) | 0 | 0 |

N = Total number of retained docked poses out of 100 with any functional site approaching the heme iron center within distance ranging between 1.5 - 6.5Å. %n = Proportion of N poses approaching the heme iron via the 4'-phenoxy spot; the major metabolic route for pyrethroids by P450 enzymes, # = Average distance between the 4'-phenoxy spot on pyrethroid poses and the heme iron in protein binding site. GBVI/WSA = Generalized-Born Volume Integral/Weighted Surface Area energy scoring function. MM = Quantum mechanics-based forcefield * = significant p-values $< 0.05$ alpha level. Thermodynamics integration simulation between insecticide and modelled proteins. ΔG (mmgb) = Binding energy (ΔG) based on single point molecular mechanics (mm) interaction energy. ΔQ (mol_charge) = molecules charge. ΔM(strain) = strain energy incurred.

**Table S4.**
Protocol for using CYP6P4b-D284E molecular diagnostic tool

| PCR Mix composition | |
|---|---|
| **Component** | **Vol X1** |
| 6P4b_ARMSOF: | 0.51 µL |
| 6P4b_ARMS_CF: | 0.51 µL |
| 6P4b_ARMS_AR: | 0.51 µL |
| 6P4b_ARMS_OR: | 0.51 µL |
| Buffer A | 1.5 µL |
| dNTP mix | 0.12 µL |
| MgCl2 | 0.75 µL |
| Polymerase (Kapa taq) | 0.12 µL |
| dH2O | 9.47 µL |

PCR Cycles

95°C - 5 min

94°C - 30 Sec

59°C - 30 sec

72°C - 1min 20 sec

(steps 94°C - 30 Sec to 72°C - 1min 20 sec: 35 cycles)

72°C - 10 min

12°C - ∞

-Make PCR reaction mix as indicated above,

-Dispense in PCR tubes,

-Add 1 µL DNA sample to be genotyped and run PCR using the following above

**Analysis**

-Run PCR products on 1.5% agarose gel

-Common band appears at 1284bp

-Resistant band appears at 527bp

-Susceptible band appears at 809bp (See Fig. 5A).

**Table S5.**
Protocol for using CYP6P4a-M220I molecular diagnostic tool

| **PCR Mix composition:** | |
|---|---|
| 1X Primetime Master Mix | 5.0 µL |
| 6p4a_F (10 µM) | 0.2 µL |
| 6p4a_R (10 µM) | 0.2 µL |
| LNA6p4a-Ile: Fam (10 µM) | 0.1 µL |
| LNA6p4a-Met: Hex (10 µM) | 0.1 µL |
| Molecular grade water | 3.4 µL |
| Genomic DNA | 1.0 µL |

PCR Cycles

95°C - 10 min ← segment 1

95°C - 10 Sec

60°C - 45 sec

72°C - 1min 20 sec

(steps 95°C - 10 Sec to 72°C - 1min 20 sec: 40 cycles (segment 2))

72°C - 10 min

12°C - ∞

**Analysis:**

-Analyse the genotypes by looking at the fluorescence for dR last.

-The FAM dye (Mutants) should be on the y-axis and HEX dye (wildtypes) on the x-axis.

-Focus on the ct values to really discriminate between genotypes (See Fig. 5B).

**Table S6.** Association between insecticide susceptibility as determined by WHO tube bioassay and CYP6P4a-M220I and CYP6P4b-D284E genotypes in An. funestus crossing between field and FANG lab colony.

| Insecticide | Comparison | OR | P value | CI |
|---|---|---|---|---|
| | | **CYP6P4b-D284E** | | |
| **Permethrin HS** | RR vs SS | 185 | P=0.001 | 8.1685 to 4189.8636 |
| | RR vs RS | 6.4 | P = 0.2315 | 0.3035 to 138.4396 |
| | RS vs SS | 39 | P = 0.0002 | 5.6823 to 267.673 |
| **Permethrin HR** | RR vs SS | 901 | P<0.0001 | 35.0574 to 23156.3417 |
| | RR vs RS | 90 | P = 0.0001 | 9.6687 to 837.7587 |
| | RS vs SS | 15.76 | P = 0.0675 | 0.8203 to 302.6730 |
| **Alphacypermethrin** | RR vs SS | 341 | P<0.0001 | 12.8969 to 9016.1915 |
| | RR vs RS | 12.19 | P = 0.0228 | 1.4161 to 104.8913 |
| | RS vs SS | 40.33 | P = 0.0126 | 2.2103 to 735.9866 |
| | | **CYP6P4a-M220I** | | |
| **Permethrin HS** | RR vs SS | 94.5 | P < 0.0001 | 12.0609 to 740.4287 |
| | RR vs RS | 2.25 | P = 0.546 | 0.1616 to 31.3308 |
| | RS vs SS | 42 | P = 0.0053 | 3.0337 to 581.4621 |
| **Permethrin HR** | RR vs SS | 885 | P <0.0001 | 34.4065 to 22763.8574 |
| | RR vs RS | 44 | P = 0.0007 | 4.9920 to 387.8210 |
| | RS vs SS | 30.39 | P = 0.0217 | 1.6472 to 560.8260 |
| **Alphacypermethrin** | RR vs SS | 189 | P = 0.0010 | 8.3163 to 4295.3047 |
| | RR vs RS | 70.4 | P = 0.1125 | 0.7304 to 19.8691 |
| | RS vs SS | 58.64 | P = 0.0055 | 3.8095 to 1039.3212 |
| | | **Genotype combination** | | |
| **Permethrin** | RR/RR vs SS/SS | 855 | P < 0.0001 | 33.2168 to 22007.6773 |
| | RR/RS vs SS/SS | 399 | P = 0.004 | 6.7856 to 23461.6639 |
| | RR/RR vs RS/RS | 70.4 | P = 0.0002 | 7.4829 to 662.3299 |
| | RR/RS vs RS/RS | 21 | P = 0.055 | 0.9306 to 473.8762 |
| | RR/RR vs RR/RS | 2.1 | P = 0.7 | 0.0720 to 63.7624 |
| | RS/RS vs SS/SS | 19 | P = 0.05 | 0.9866 to 365.8996 |
| **Alphacypermethrin** | RR/RR vs SS/SS | 172,2 | P = 0.0012 | 7.5569 to 3923.9444 |
| | RR/RS vs SS/SS | 90 | P = 0.0055 | 3.7550 to 2166.7357 |
| | RR/RR vs RS/RS | 4.7 | P = 0.0704 | 0.8792 to 24.9923 |
| | RR/RS vs RS/RS | 4.7 | P = 0.1802 | 0.4893 to 44.9061 |
| | RR/RR vs RR/RS | 1 | P = 1.000 | 0.0721 to 13.8684 |
| | RS/RS vs SS/SS | 43.7 | P = 0.0104 | 2.4260 to 785.2074 |

**Table S7.** Association between insecticide susceptibility as determined by WHO cone bioassay and CYP6P4a-M220I and CYP6P4b-D284E genotypes in An. funestus crossing between field and FANG lab colony.

| Bed net | Comparison | OR | P value | CI |
|---|---|---|---|---|
| | | **CYP6P4b-D284E** | | |
| **Permanet 2.0** | RR vs SS | 5.7273 | P = 0.0111 | 1.4901 to 22.0124 |
| | RR vs RS | 1.0588 | P = 0.9391 | 0.2446 to 4.5830 |
| | RS vs SS | 5.4091 | P = 0.0051 | 1.6579 to 17.6481 |
| **Olyset** | RR vs SS | 35.7778 | P<0.0001 | 6.3253 to 202.3699 |
| | RR vs RS | 2.5667 | P = 0.2019 | 0.6034 to 10.9184 |
| | RS vs SS | 13.95 | P = 0.0003 | 3.4023 to 57.1108 |
| **Duranet** | RR vs SS | 19.06 | P<0.0001 | 6.4130 to 56.6413 |
| | RR vs RS | 2.488 | P = 0.0126 | 1.241 to 4.787 |
| | RS vs SS | 7.66 | P<0.0001 | 2.7712 to 21.1707 |
| | | **CYP6P4a-M220I** | | |
| **Permanet 2.0** | RR vs SS | 6.682 | P = 0.0205 | 1.360 to 34.35 |
| | RR vs RS | 1.4318 | P=0.6881 | 0.2594 to 8.6736 |
| | RS vs SS | 4.67 | P = 0.0054 | 1.6560 to 14.4989 |
| **Olyset** | RR vs SS | 27 | P<0.0001 | 4.431 to 141.3 |
| | RR vs RS | 1.63 | P=0.5736 | 0.3591 to 8.516 |
| | RS vs SS | 16.57 | P<0.0001 | 4.972 to 49.57 |
| **Duranet** | RR vs SS | 10.5 | P<0.01 | 2.045 to 45.56 |
| | RR vs RS | 2.444 | P=0.218 | 0.6379 to 9.072 |
| | RS vs SS | 4.295 | P<0.05 | 1.289 to 13.06 |
| | | **Genotype combination** | | |
| **Permanet 2.0** | RR/RR vs SS/SS | 6.6818 | P = 0.0317 | 1.1816 to 37.7865 |
| | RR/RR vs RS/RS | 1.3125 | P = 0.7709 | 0.2105 to 8.1844 |
| | RR/RR vs RR/RS | 1.167 | P = 0.892 | 0.1238 to 10.9909 |
| | RS/RS vs SS/SS | 5.0909 | P = 0.0073 | 1.5511 to 16.7092 |
| | RR/RS vs SS/SS | 5.7273 | P = 0.0518 | 0.9866 to 33.2482 |
| | RR/RS vs RS/RS | 1.125 | P = 0.9 | 0.1760 to 7.1915 |
| **Olyset** | RR/RR vs SS/SS | 38.5 | P = 0.0008 | 4.5452 to 326.1145 |
| | RR/RR vs RS/RS | 1.5556 | P = 0.6263 | 0.2627 to 9.2108 |
| | RS/RS vs SS/SS | 24.75 | P = 0.0002 | 4.6589 to 131.4817 |
| | RR/RR vs RR/RS | 0.438 | P = 0.534 | 0.0323 to 5.9260 |
| | RR/RS vs SS/SS | 88 | P = 0.0005 | 6.9874 to 1108.2746 |
| | RR/RS vs RS/RS | 3.5556 | P = 0.2669 | 0.3787 to 33.3823 |
| | RR/RR vs RS/SS | 14 | P = 0.055 | 0.9441 to 207.6072 |
| | RS/RS vs RS/SS | 4.5 | P = 0.12 | 0.6793 to 29.8086 |
| | RR/RS vs RS/SS | 20 | P = 0.03 | 1.3908 to 287.6143 |
| **Duranet** | RR/RR vs SS/SS | 10 | P = 0.0089 | 1.7809 to 56.1515 |
| | RR/RR vs RS/RS | 2.5185 | P = 0.2223 | 0.5714 to 11.1001 |
| | RR/RR vs RR/RS | 0.8 | P = 0.813 | 0.1257 to 5.0919 |
| | RS/RS vs SS/SS | 3.9706 | P = 0.0357 | 1.0965 to 14.3785 |
| | RR/RS vs SS/SS | 12.5 | P = 0.0035 | 2.2895 to 68.2473 |
| | RR/RS vs RS/RS | 3.1481 | P=0.12 | 0.7381 to 13.4283 |

#### Table S8. Primers used in the study

| Primer ID | Forward primer | Reverse primer |
|---|---|---|
| | **Full gene amplification** | |
| CYP6P4a_full | ATGGATTTTTTGGGCTATGTGTTG | CCTTTTACACCCACCAGGAA |
| CYP6P4b_full | ATGGATCTTCTGGGTTATGTGTTG | GTATGTCCGTTCTGCACCC |
| | **Cloning for in vitro heterologous expression** | |
| ompA+2 F | GGCCGGCCATATGAAAAAGACAGCTATCGCG | |
| ompA+2_6P4aR linker | CAACACATAGCCCAAAAAATCCATCGGAGCGGCCTGCGCTACGGTAGCGAA | |
| ompA_6P4a_R | TCTAGAGTCGACTTAAAGCTTATCAATCTTTAG | |
| OMPA+2_6P4b R linker | CAACACATAACCCAGAAGATCCATCGGAGCGGCCTGCGCTACGGTAGCGAA | |
| ompA_6P4b_R | GTCGACTCTAGATCAGAAACCTTCAATCTTATCAAC | |
| | **Transgenic flies** | |
| CYP6P4a_pUASattB | CGGCCGATGGATTTTTTGGGCTATGTGTTG | TCTAGATTAATATCAATCTTTAGATAATTTCC |
| CYP6P4b_pUASattB | CGGCCGATGGATCTTCTGGGTTATGTGTTG | TCTAGATCAGAAACCTTCAATCTTATCTACC |
| | **Fly qPCR** | |
| 6P4B_ Fly_qPCR | GGTCCACGGATTTGTATTGG | ATACGAGTTCGGACGGTGTT |
| 6P4A_ Fly_qPCR | AGCTTACGCCAACGTTTACC | GTCCCGATCACATCCGTAGT |
| RPL11 | CGATCCCTCCATCGGTATCT | AACCACTTCATGGCATCCTC |
| | **CYP6P4a-M2201molecular marker** | |
| 6p4a_F: | ATACGGCAACAAGGTGTTC | CCTTCGTCAGTCAGCTTAAC |
| LNA6p4a-Met:Hex | TGT+TCTTA+T+G+GT+AA+A+GT | |
| LNA6p4a-Ile:Fam | ACTGT+C+CTTAT+T+TT+C+AA+AT | |
| | **CYP6P4b-D284E molecular marker** | |
| 6P4b_ARMSOF | CTATCTGCTCATCTGTTTGCACTGGA | |
| 6P4b_ARMSOR | TGACCATTTCATATGTCACTTGTCCAC | |
| 6P4b_ARMS_CF | CTGCAGATTAAGAACAAAGGTTATTTGAAC | |
| 6P4b_ARMS_AR | TTATCATTGGCTCCAATGTCACGTTTTT | |
| | **Primers used for qRT-PCR gene quantification** | |
| CYP6P4a-qPCR | AACTCGTATTCGACCCCAAA | CGTTTCCATGGAATTACATTTTCT |
| RSP7 | GTGTTCGGTTCCAAGGTGAT | TCCGAGTTCATTTCCAGCTC |
| ACTIN | TTAAACCCAAAAGCCAATCG | ACCGGATGCATACAGTGACA |
